# Ventral Pallidum GABA Neurons Mediate Motivation Underlying Risky Choice

**DOI:** 10.1101/2020.08.04.221960

**Authors:** M.R. Farrell, J.S.D. Esteban, L. Faget, S.B. Floresco, T.S. Hnasko, S.V. Mahler

**Author notes:** **Corresponding Author**: Mitchell R. Farrell, M.S., Department of Neurobiology & Behavior. University of California, Irvine, 1132A McGaugh Hall, Irvine, CA 92697, (O).

## Abstract

Pursuing rewards while avoiding danger is an essential function of any nervous system. Here, we examine a new mechanism helping rats negotiate the balance between risk and reward when making high-stakes decisions. Specifically, we focus on GABA neurons within an emerging mesolimbic circuit nexus—the ventral pallidum (VP). These neurons play a distinct role from other VP neurons in simple motivated behaviors in mice, but their roles in more complex motivated behaviors is unknown. Here, we interrogate the behavioral functions of VP^GABA^ neurons in male and female transgenic GAD1:Cre rats (and wildtype littermates), using a reversible chemogenetic inhibition approach. Employing a behavioral assay of risky decision making, and of the food-seeking and shock-avoidance components of this task, we show that engaging inhibitory G_i/o_ signaling specifically in VP^GABA^ neurons suppresses motivation to pursue highly salient palatable foods, and possibly also motivation to avoid being shocked. In contrast, inhibiting these neurons did not affect seeking of low-value food, free consumption of palatable food, or unconditioned affective responses to shock. Accordingly, when rats considered whether to pursue food despite potential for shock in a risky decision-making task, inhibiting VP^GABA^ neurons caused them to more readily select a small but safe reward over a large but dangerous one—an effect not seen in the absence of shock threat. Together, results indicate that VP^GABA^ neurons are critical for high-stakes adaptive responding that is necessary for survival, but which may also malfunction in psychiatric disorders.

**Significance Statement:** In a dynamic world, it is essential to implement appropriate behaviors under circumstances involving rewards, threats, or both. Here, we demonstrate a crucial role for VP^GABA^ neurons in high-stakes motivated behavior of several types. We show that this VP^GABA^ role in motivation impacts decision making, as inhibiting these neurons yields a conservative, risk-averse strategy not seen when the task is performed without threat of shock. These new roles for VP^GABA^ neurons in behavior may inform future strategies for treating addiction, and other disorders of maladaptive decision making.

## Introduction

Executing appropriate action under conflicting motivations is fundamental for survival in a dynamic world. For example, balancing appetitive and aversive motivations is essential for most animals to eat without being eaten. In humans, this interplay of motivations is required for appropriate decision making, and inappropriately balancing reward and aversion likely contributes to a variety of psychiatric disorders including addiction. Indeed, compulsive drug use and relapse in addiction can be conceptualized as desire for drugs overcoming the perceived threat of consequences, leading to poor decisions. Yet most preclinical studies explore reward in the absence of threat, or threat without reward—conditions that rarely occur in the lives of opportunistic prey species like rodents. Understanding how functionally distinct cell populations within brain motivation circuits participate in appetitive, aversive, and also *mixed motivations* will provide novel insights into the neural substrates of both adaptive and maladaptive decision making.

The ventral pallidum (VP) is at an anatomical interface of motivation and action (Heimer et al., 1982), and is ideally positioned to contribute to behavioral responses to both rewards and threats. Across species, VP neurons encode the motivational value of specific actions that result in reward, in a manner that reflects whether such actions are worth generating (Pessiglione et al., 2007; Tindell et al., 2009; Tachibana and Hikosaka, 2012; Richard et al., 2016; Fujimoto et al., 2019). VP also plays a causal role in reward, as pharmacological stimulation enhances spontaneous food intake (Stratford et al., 1999; Smith et al., 2009) and hedonic evaluations of tastes (Berridge and Kringelbach, 2015), whereas perturbing VP disrupts conditioned motivation (McAlonan et al., 1993; Chang et al., 2015), and reward-related working memory (Floresco et al., 1999). Notably, VP also plays a crucial role in seeking of multiple classes of addictive drugs (Rogers et al., 2008; Mahler et al., 2014; Farrell et al., 2019; Heinsbroek et al., 2019; Prasad and McNally, 2020).

However, it has become clear that VP not only contributes to reward, but also to aversive motivational processes. Pharmacological disinhibition of VP neurons generates spontaneous defensive behavior in rats (Smith and Berridge, 2005), and disrupts the ability of monkeys to avoid a cued aversive airpuff (Saga et al., 2016). Perhaps relevant to this are recent reports revealing that a glutamatergic subpopulation of VP neurons mediates aversive motivation and learning in mice, as they fire in response to aversive stimuli, promote avoidance and curtail reward seeking when optogenetically stimulated, and generally cause opposite effects when optogenetically inhibited (Faget et al., 2018; Tooley et al., 2018; Heinsbroek et al., 2019; Stephenson-Jones et al., 2020).

In contrast to VP glutamate neurons, VP^GABA^ neurons have instead been linked to reward seeking and approach responses in mice. For example, photostimulating VP^GABA^ neurons is reinforcing, and induces food intake (Zhu et al., 2017; Faget et al., 2018; Stephenson-Jones et al., 2020). VP^GABA^ neurons also selectively fire to reward cues, and their activity is required for operant reward seeking, but not avoidance responses (Stephenson-Jones et al., 2020). These results support the notion of extensive functional heterogeneity amongst VP cell populations (Smith and Berridge, 2005; Kupchik and Kalivas, 2013; Mahler et al., 2014; Root et al., 2015), and show that VP^GABA^ neurons play a distinct, though poorly characterized role in behavior.

Here we systematically characterize the behavioral functions of VP^GABA^ neurons, in transgenic GAD1:Cre rats. Using validated, specific, and reversable chemogenetic inhibition of VP^GABA^ neurons, we show they mediate both highly motivated pursuit of salient foods, and avoidance of shocks. In contrast, inhibiting these cells does not affect shock-induced aversion, low-motivation food seeking, free food consumption, or locomotion. Notably, when rats made choices about food rewards under threat of shock, VP^GABA^ inhibition shifted choice bias towards a more risk averse strategy, increasing preference for small/safe rewards over large/risky ones. Together, these results show that VP^GABA^ neurons govern high-stakes motivational processes underlying risky decision making.

## Methods

### Subjects

Male (*n* = 46) and female (*n* = 35) Long-Evans hemizygous GAD1:Cre rats (Sharpe et al., 2017; Gibson et al., 2018; Wakabayashi et al., 2019) and their Cre-negative wildtype littermates (WT) were pair-housed in polycarbonate tub-style cages (48 × 20 × 27 cm) with bedding and nesting material. Rats were maintained on a reverse 12 hr light-dark cycle, with testing in the dark phase. Water was available *ad libitum* and food was restricted to ∼90% of free-feeding weight during behavioral testing (∼6-9 g/day/rat), unless otherwise noted. During food restriction, food was placed in the homecage after each behavioral testing session. The cohorts of rats used for each of the behavioral tasks, the sex distribution of each cohort, and the chronological ordering of behavioral testing is presented in **Table 1**. All procedures were approved by the University of California Irvine Institutional Animal Care and Use Committee, and are in accordance with the NIH Guide for the Care and Use of Laboratory Animals.

**Table 1.** Chronological order of behavioral testing for each cohort of rats. Chronological order of behavioral testing for each cohort of rats organized left (first test) to right (last test), with each behavioral test indicated on the top row. Cohort sex breakdown (M=male, F=female): Cohort 1 (7M, 8F), 2 (11M, 5F), 3 (5M, 11F), 4 (4M, 4F), 5 (8M, 0F), 6 (11M, 7F).

| Risky Decision Task | Progressive Ratio | FR1 Palatable (hungry/sated) | Avoidance/Escape Task | Motor Reactivity | Ultrasonic Vocalizations | Reward Magnitude Discrimination | FR1 Chow (hungry/sated) | M&M Free Access | Locomotion |
| --- | --- | --- | --- | --- | --- | --- | --- | --- | --- |
| Cohort 1 | Cohort 1 |  |  |  |  |  |  |  |  |
| Cohort 2 | Cohort 2 |  | Cohort 2 | Cohort 2 |  |  |  |  |  |
| Cohort 3 | Cohort 3 | Cohort 3 | Cohort 3 |  | Cohort 3 |  |  |  |  |
|  | Cohort 4 |  |  |  |  |  |  |  |  |
|  |  |  |  |  |  | Cohort 5 | Cohort 5 |  |  |
|  |  |  |  |  |  |  |  | Cohort 6 | Cohort 6 |

### Chemogenetic Methods

#### Surgery and Viral Vectors

Rats were anesthetized with ketamine (56.5 mg/kg) and xylazine (8.7 mg/kg), and treated for pain with meloxicam (1.0 mg/kg). An adeno-associated vector containing double-floxed, inverted open reading frame (DIO) mCherry-tagged hM4Di designer receptors (Armbruster et al., 2007) (DREADDs; AAV2-hSyn-DIO-hM4D(Gi)-mCherry; titer: 1 × 10^12^ GC/mL; Addgene) was injected bilaterally into VP (relative to bregma: AP 0.0 mm, ML ±2.0 mm, DV -8.2 mm; ∼300 nL/hemisphere) using a Picospritzer and glass micropipette. Injections occurred over 1 min, and the pipette was left in place for 5 min after injection to limit spread. Both GAD1:Cre and WT rats were injected with the active hM4Di DREADD virus, and lack of hM4Di/mCherry expression was confirmed in each WT rat.

#### Drugs

Clozapine-N-oxide (CNO) was obtained from NIDA, and subsequently stored at 4° C in powder aliquots stored in desiccant, and protected from light. CNO was dissolved in a vehicle containing 5% dimethyl sulfoxide (DMSO) in saline, and injected at 5 mg/kg intraperitoneally (IP), 30 min prior to tests. For microinjections, bicuculline methiodide (Sigma) was dissolved in artificial cerebrospinal fluid (Thermofisher), stored in aliquots at -20° C, and thawed just prior to use.

### DREADD Validation

#### Localization of DREADD Expression to VP

Virus expression in GAD1:Cre rats was amplified with mCherry immunohistochemistry, and sections were co-stained for substance P, an anatomical marker of VP borders. First, behaviorally-tested rats were perfused with cold 0.9% saline and 4% paraformaldehyde after completion of experiments. Brains were cryoprotected in 20% sucrose, sectioned at 40 μm, and blocked in 3% normal donkey serum PBST. Tissue was incubated 16 hrs in rabbit anti-substance P (ImmunoStar; 1:5000) and mouse anti-mCherry antibodies (Clontech; 1:2000) in PBST-azide with 3% normal donkey serum. After washing, slices were incubated in the dark for 4 hrs in Alexafluor donkey anti-Rabbit 488 and donkey anti-Mouse 594 (Thermofisher), then washed, mounted, and coverslipped with Fluoromount (Thermofisher). mCherry expression was imaged at 10x, and the zone of expression in each hemisphere of each rat was mapped in relation to VP borders, and a rat brain atlas (Paxinos and Watson, 2006).

#### Localization of DREADDs Specifically to VP^GABA^ Neurons

Experimentally-naïve GAD1:Cre rats (*n* = 4) injected in VP with AAV2-hSyn-DIO-mCherry were euthanized, and fresh brains were immediately extracted and frozen in isopentane before storage at −80 °C. Brains were serially cut (20 μm) on a cryostat, and placed directly onto slides before returning to storage at −80 °C. Three different coronal sections of the VP near the center of mCherry expression were used per brain. In situ hybridizations were performed using the RNAscope Multiplex Fluorescent Assay (Advanced Cell Diagnostics). RNA hybridization probes included antisense probes against rat *Gad1* (316401-C1), rat *Slc17a6* (vglut2 gene; 317011-C3) and *mCherry* (431201-C2) (*n* = 2), or antisense probes against rat *Gad1* (316401-C1), rat *Slc32a1* (vgat gene; 424541-C3) and *mCherry* (431201-C2) (*n* = 2), both respectively labeled with alexa488, atto647, and atto550 fluorophores. DAPI was used to label nuclei and identify cells. Three images/hemisphere/section were taken at 63x (1.4 NA) magnification using a Zeiss AxioObserver Z1 widefield Epifluorescence microscope with a Zeiss ApoTome 2.0 for structured illumination and Zen Blue software for counting. Wide-field images were taken at 20x (0.75 NA) magnification. Cells that exhibited at least 4 puncta (RNA molecules) in addition to DAPI were counted as expressing the respective gene.

#### DREADD-Dependent Inhibition of VP^GABA^ Neurons by CNO

In order to verify the ability of CNO to inhibit VP neurons in a DREADD-dependent manner, we tested the ability of systemic CNO to inhibit exogenously-stimulated VP neural activity. Experimentally-naïve GAD1:Cre rats (*n* = 3) were injected unilaterally with the previously described AAV2 DIO-hM4Di-mCherry vector in ipsilateral VP, and contralaterally in VP with a matched AAV2 DIO-mCherry control vector (4.7 × 10^12^ GC/mL, AddGene). Three weeks later, bilateral intracranial cannulae were implanted 2 mm dorsal to the injection target, using previously described procedures (Mahler et al., 2013a; Mahler et al., 2014; Mahler et al., 2019), and rats recovered for 5 d. Rats were then injected systemically with CNO, in order to engage unilaterally expressed VP^GABA^ hM4Di receptors. 30 min later, rats were bilaterally injected in VP with 0.5 µL of bicuculline (0.01 μg/0.5 μL/50 sec), inducing neural activity in the local VP area in both hemispheres. 90 min later, rats were perfused, and brains were processed for Fos and mCherry to determine whether bicuculine-induced Fos was suppressed by hM4Di activation (i.e. if there was less Fos expression in the hM4Di hemisphere than the mCherry hemisphere). VP sections near the center of the microinjection sites were incubated overnight at room temperature in rabbit anti-Fos (1:5000; Millipore) and mouse anti-DSRed (targeting mCherry; 1:2000; Clontech), washed, incubated in Alexafluor donkey anti-Rabbit 488 and donkey anti-Mouse 594 in dark for 4 hrs at room temperature, then coverslipped as above. For each rat, 2-3 brain sections/hemisphere/rat with VP-localized microinjector tip damage were selected for manual quantification at 10x magnification by an observer blind to experimental manipulation. mCherry-only, and mCherry/Fos co-expressing cells within VP borders (Paxinos and Watson, 2006) were counted in both hemispheres. The percentage of mCherry cells co-expressing Fos in each sample was calculated, and per-hemisphere averages were computed for each rat for statistical analysis.

### Behavioral Testing Methods

#### Risky Decision-Making Task

##### Operant Apparatus

All operant testing was performed in Med Associates operant chambers in sound-attenuating boxes, equipped with two retractable levers with associated stimulus lights above them. Between the two levers was a food magazine connected to a food pellet dispenser. Two nose-poke ports were positioned on the opposite wall with a yellow light in one of the ports. Boxes were equipped with tone/white noise and footshock generators.

##### Habituation Training

We adapted a previously-reported risky decision task and associated training protocol (Simon et al., 2009; Simon and Setlow, 2012; Orsini et al., 2015a). Mildly food deprived male (*n* = 23) and female rats (*n* = 22) were familiarized to highly palatable, banana-flavored, sucrose, fat, and protein-containing pellets in their homecage (Bio-Serv, Ct # F0024), then on day 1 of training, 38 pellets were delivered into the food magazine on a variable time 100 sec schedule (140 sec, 100 sec, 60 sec) during a single ∼60 min session. Rats that failed to eat >19 pellets were given a second day of magazine training.

##### Lever Pressing Training

Next, rats were trained to lever press for the banana pellets in daily 30 min sessions. Each session began with illumination of the house light, and extension of a single lever plus illumination of the associated stimulus light (right or left, counterbalanced). One pellet and a brief auditory tone cue (0.5 sec, 2.9 kHz) were delivered on a fixed ratio 1 (FR1) schedule, with a 10 sec timeout period between pellet deliveries. Daily FR1 training continued until criterion was met (50 pellet/30 min session), followed by training on the alternate (left or right) lever, again until criterion. *Lever Choice Training.* The next training phase consisted of daily 1 hr sessions that taught rats to press levers within 10 sec of their extension. Sessions began with illumination of the houselight, and every 40 sec one lever (right or left) was extended for 10 sec, along with the associated stimulus light. Lever presses yielded 1 food pellet, and the tone cue. If no press occurred during the 10 sec extension window, the lever retracted and stimulus light extinguished, the trial was counted as an omission, and rats were required to wait until the next lever extension trial. Each session consisted of 35 left lever, and 35 right lever extensions with a 40 sec intertrial interval, independent of the rats’ pressing or omitting. Rats that met criterion (<10 omissions) on two consecutive sessions were moved to the next phase of the task. In this phase procedures were the same, except that now pressing one lever (left or right, counterbalanced) delivered 1 pellet accompanied by the tone cue, and pressing the other lever delivered 2 pellets and 2 tone cues. Rats were trained for at least 3 d in this manner, until 2 consecutive days with <10 omissions.

##### Risky Decision Task

Rats were next trained on the risk task, in which the threat of shock was introduced. At session start, as above one lever yielded 1 pellet, and the other 2 throughout the session. However, now the 2-pellet option came with the chance of concurrently-delivered shock; the probability of which increased over the course of the session. Sessions consisted of 5 blocks with 20 trials each, for a total of 66 min. Blocks represent changes in footshock probability associated with large/risky lever presses such that in the first 20-trial block there was no chance of shock, and in each subsequent block shock probability increased by 25% (Block 1: 0% probability, Block 2: 25%, Block 3: 50%, Block 4: 75%, Block 5: 100%). Each 20-trial block began with 8 ‘forced choice’ trials in which a single lever was extended (4 large/risky and 4 small/safe lever extensions, random order) to establish the shock contingency for that block. Following the 8 forced choice trials, 12 ‘free choice’ trials commenced in which both the large/risky and small/safe levers were extended simultaneously to allow choice of the preferred option (small/safe; large/risky). If no lever press occurred within 10 sec, the lever(s) were retracted, stimulus light(s) extinguished, and the trial was considered an omission. Footshock intensity (mA) was titrated individually for each rat to ensure sufficient parametric space to observe either increases or decreases in risky choice, as reported previously (Orsini et al., 2017). Footshock intensity started at 0.15 mA for each rat upon beginning the risky decision task, and percent choice of the large/risky reward was monitored daily for fluctuations in decision making. Footshock intensity was increased or decreased each day by 0.05 mA, until stable decision making behavior was achieved in all animals.

##### Stable Pre-Test Baseline Performance

Rats generally achieved stability within 10-20 sessions, with near-exclusive choice of the “risky” 2 pellet option when chance of shock was zero, then a parametric shift to the “safe” 1 pellet option as shock probability increased across blocks (interaction of block X lever: *F*_(4, 132)_ = 111.5, *p* < 0.0001). Rats were trained until performance was stable for 5 consecutive days (no difference in 5 d average performance pre-vehicle/CNO: *F*_(1, 229)_ = 0.45, n.s.; or interaction of block x treatment: *F*_(4, 229)_ = 0.023, n.s.), then were assigned to receive counterbalanced vehicle and CNO tests, between which behavior was re-stabilized over ∼5 days of training. Over the course of these experiments, 6 rats made >50% omissions during vehicle treatment sessions (range 50-72 omissions over 100 trials), making interpretation of their data problematic. Accordingly, their data were excluded from risky decision analyses.

#### Reward Magnitude Discrimination

To characterize potential VP inhibition effects on mere preference for larger versus smaller rewards, a separate cohort of GAD1:Cre rats (*n* = 8) were trained identically to above, except that shock was never introduced. We then evaluated CNO effects (versus vehicle) on choice of the 2-pellet lever over the 1-pellet lever, in the absence of shock. Rats required ∼5-10 training sessions before displaying stable preference for the larger reward, after which they received CNO and vehicle tests on separate days.

After the first test, rats received ∼3 days of training to re-stabilize performance, then were given their second counterbalanced treatment.

#### Spontaneous Palatable Food Intake

*Ad libitum*-fed rats (*n* = 18) were placed in polycarbonate cages (44.5 x 24 x 20 cm) with bedding and ∼12 g of peanut butter M&M chocolates for 1 hr on 2 consecutive days, to habituate them to test conditions. The next day, rats were administered CNO or vehicle (counterbalanced, separate days) 30 min prior to a 1 hr intake test. 48 hrs later the procedure was repeated with the other drug treatment. Food intake (g) was measured.

#### High Effort Instrumental Responding for Palatable Food

To assess the involvement of the VP in food seeking under higher effort requirements, mildly food-deprived rats (*n* = 39) were trained to nosepoke on a progressive ratio schedule of reinforcement. Sessions began with illumination of the both the houselight and a light within the active nosepoke port. When the required schedule was achieved, 3 banana pellets + 3 concurrently-delivered 0.5 sec white noise pulses were delivered. The number of nosepokes required for reward increased each time the prior requirement was achieved (FR 1, 6, 15, 20, 25, 32, 40, 50, 62, 77, 95, 118, 145, 178, 219, 268, 328, 402, 492, 603 (Smith and Aston-Jones, 2012)). Inactive port entries were inconsequential, but recorded. Sessions lasted a maximum of 2 hrs, or less if the rat failed to reach the next ratio within 20 min of achieving the prior one. Training continued until rats achieved stable performance for 2 consecutive sessions (<25% change in active nosepokes). Pressing was re-stabilized between counterbalanced vehicle/CNO tests.

#### Low Effort Instrumental Responding for Palatable Food or Conventional Chow

Mildly-food deprived rats or *ad libitum* fed rats (*n* = 16) were trained to nosepoke for palatable 45 mg banana pellets on an FR1 schedule of reinforcement during daily 1 hr sessions. Separate animals (*n* = 8) were trained to respond for 45 mg chow pellets (Bio-Serv, Ct # F0165) instead using the same procedures. Sessions began with illumination of the houselight and active nosepoke port light. Active nosepokes resulted in delivery of a pellet into the food cup, while inactive nosepokes were without consequence. Rats were trained until achieving stability (<25% change in active nosepokes) for 2 consecutive sessions (2-7 sessions), then tested with counterbalanced vehicle/CNO, with 1+ days of re-stabilization between tests.

#### Operant Shock Avoidance/Escape Task

Procedures were adapted from a previously described shock avoidance/escape task (Oleson et al., 2012). Rats (*n* = 18 trained) that had previously performed the risky decision task, progressive ratio task, and palatable food FR1 task were tested, and footshock intensity (mA) was the same as that used for the rat during the previously-trained risk task (0.15-45 mA). Each 30 min session began with illumination of the houselight, and every 20 sec an active and inactive lever were extended. Initial training taught rats to press a lever to turn off a repeated foot shock. During this initial ‘escape only’ training, lever extension was met with a concurrent footshock that repeated (0.1 sec footshock every 2 sec) until the active lever was pressed, at which time footshock ceased, both active and inactive levers were retracted, and a 20 sec white noise safety signal was played. Then the next trial began with re-extension of both levers, a sequence that was repeated until the end of the 30 min session. Training proceeded for at least 2 d, until consistent escape behavior was observed.

Next, rats were trained to avoid, as well as to escape shocks in 30 min sessions. In this phase, levers were again extended at the start of each trial, but now this occurred 2 sec prior to initiation of shocks. If the active lever was pressed in this 2 sec period (an avoid response), no shock occurred, levers retracted, and the safety signal was played for 20 sec. If no press occurred before 2 sec elapsed, repeating footshock commenced as above, until an active lever press occurred (escape response), at which time levers were retracted and the safety signal was played for 20 sec. Inactive lever presses were inconsequential but recorded. All rats with >5 avoidance lever presses on the vehicle test day were included for analyses (*n* = 18). Rats were administered counterbalanced vehicle and CNO tests 30 min prior to avoidance/escape sessions, with ∼3 d between tests to re-stabilize behavior. Data were analyzed by assessing 1) the change in the ratio of avoidance presses to escape presses from pre-test baseline (i.e. change from baseline avoidance %), 2) the ratio of avoidance presses to escape presses on vehicle and CNO tests (i.e. raw avoidance %), 3) latency to avoid footshock, and 4) latency to escape repeated footshock.

#### Motor Responses to Shock

To query the role of VP^GABA^ in affective responses to shock itself, rats (*n* = 16) were tested for overt motor reactions to shocks of ascending intensity in a chamber in which they had not been previously tested. The houselight was illuminated and 2 min elapsed. This waiting period ended with one 0.30 mA footshock to limit ongoing exploration. Following this shock, rats were administered 5 consecutive 1 sec, 0.05 mA shocks, each separated by 10 sec. After these 5 shocks, the procedure was repeated with blocks of increasingly intense shocks, increasing by 0.05 mA with each block. Motor reactivity was evaluated during testing, according to previously published criteria (Bonnet and Peterson, 1975). Briefly, motor reactivity was separated into 4 categories: 0: no movement, 1: flinch of a paw or a startle response, 2: elevation of one or two paws, 3: rapid movement of three or all paws. When 3 out of 5 shocks at a particular intensity elicited level 3+ motor reactivity, the session was terminated. CNO/vehicle tests were counterbalanced, and administered 48 hrs apart.

#### Ultrasonic Vocalization Responses to Shock

To further query affective shock responses, rats (*n* = 16) were administered 2 shock-induced ultrasonic vocalization tests after counterbalanced vehicle or CNO, held 48 hrs apart. Recordings again occurred in a chamber in which they had not been previously tested. Sessions began with illumination of the houselight, and following a 2 min baseline period, rats received 5 unsignaled footshocks (1 sec, 0.75 mA), each separated by 1 min. Recordings were made with condenser ultrasound microphones (frequency range: 10–200 kHz; CM16/CMPA, Avisoft Bioacoustics, Berlin, Germany) that were centered atop the operant chamber and pointed directly toward the center of the chamber (∼18 cm above the floor). USV recordings were made on an UltraSoundGate 416H data acquisition device (Avisoft Bioacoustics; sampling rate 250 kHz; 16-bit resolution), as reported previously (Mahler et al., 2013b).

Spectrograms were visualized using Avisoft software, and ultrasonic vocalizations (USVs) were manually quantified by an observer blind to experimental conditions. Aversion-related 22 kHz USVs were operationalized as 18-30 kHz with a duration greater than 10 ms, and positive affect-related high frequency USVs were operationalized as those >30 kHz frequency, with a duration greater than 10 ms.

#### General Locomotor Activity

General locomotor activity was assessed in a locomotor testing chamber (43 × 43 × 30.5?cm) with corncob bedding. Following two daily 2 hr habituation sessions, infrared beams captured horizontal distance traveled and number of vertical rears following vehicle/CNO injections (counterbalanced tests, 48 hrs apart).

#### Statistics and Analyses

Graphpad Prism and SPSS were used for all statistical analyses. CNO and vehicle tests were counterbalanced for each experiment. An independent samples *t*-test was used to compare %Fos in ipsilateral mCherry+ VP neurons versus %Fos in contralateral hM4Di-mCherry neurons following bicuculline microinjection and systemic CNO injection. Male and female footshock intensities required on the risky decision task were compared with an independent samples *t*-test. For the risky decision task, reward magnitude discrimination, and motor shock reactivity tasks, effects of drug and block were analyzed with separate two-way ANOVAs in GAD1:Cre and WT rats, along with Sidak posthoc tests. Win-stay and lose-shift behavior was characterized on choice trials, and analyzed with separate paired sample *t*-tests for GAD1:Cre and WT rats. Win-stay was operationalized as the number of risky choices after a non-shocked risky choice, divided by the total number of non-shocked risky choices, whereas lose-shift was the number of safe choices followed by a shocked risky choice, divided by the total number of shocked risky choices. In addition, we performed two-way ANOVAs with treatment (vehicle and CNO) and genotype (WT and GAD1:Cre) as factors for relevant comparisons, along with a third factor (bin) for risk task data. Pearson’s correlation was used to determine whether footshock intensity employed in the risky decision task correlated with pressing for the large/risky over the small/safe option. Separate one-way ANOVAs were conducted to ask whether latency to press the small/safe or large/risky options increased across the session when tested with vehicle treatment. Latency data for the risky decision task excluded all trials in which omissions occurred. For avoidance/escape task, FR1, and progressive ratio tasks, effects of CNO versus vehicle in GAD1:Cre and WT rats were analyzed with paired sample t-tests. Due to the high variability in USV production among rats, all USV data were analyzed as percent of vehicle test day, and compared to 100% with one-sample *t*-test. Sample sizes were chosen based on those used in prior experiments (Simon and Setlow, 2012; Orsini et al., 2017; Farrell et al., 2019). Two-tailed tests with a significance threshold of *p* < 0.05 were used for all analyses.

## Results

### Selective, Functional hM4Di Expression in VP^GABA^ Neurons

GAD1:Cre rats exhibited hM4Di-mCherry expression (*n* = 53) that was largely localized within substance P-defined VP borders (**Fig 1A-B**). Specifically, at the center of expression, mean ± SEM = 68.9% ± 1.2 of total viral expression area was localized within VP, and in behaviorally tested rats 68.8% ± 1.3 of VP area contained mCherry expression. At least 55% of DREADD expressing cells were localized within VP borders of included rats, though most animals also had at least some expression in adjacent GABAergic structures. At least some extra-VP expression was observed in lateral preoptic area of 20.8% of rats, bed nucleus of the stria terminalis (30.2%), horizontal limb of the diagonal band of Broca (66%), and globus pallidus (15.1% of rats). Verifying specificity of expression, we used RNAscope to show that mCherry mRNA was largely colocalized with GABA-specific markers *gad1* and *vgat* mRNA in VP (mCherry+gad1+: m = 91.28 ± 2.6; mCherry+vgat+: m = 91.58 ± 2.75) (data not shown). The vast majority of these neurons were triple labeled for *mCherry, gad1*, and *vgat* (mCherry+gad1+vgat+: m = 87.92 ± 2.58), indicating robust expression in GABAergic VP neurons. Little mCherry expression was detected in vglut2+ neurons (mCherry+vglut2+: m = 8.42 ± 3.42), and of these mCherry+vglut2+ cells, 40.6% also localized with *gad1* (mCherry+vglut2+gad1+: m = 3.42 ± 1.08) (**Fig 1C-E**), possibly indicating co-expression of GABA and glutamate in some pallidal neurons (Meye et al., 2016; Faget et al., 2018; Farrell et al., 2019).

**Figure 1.**
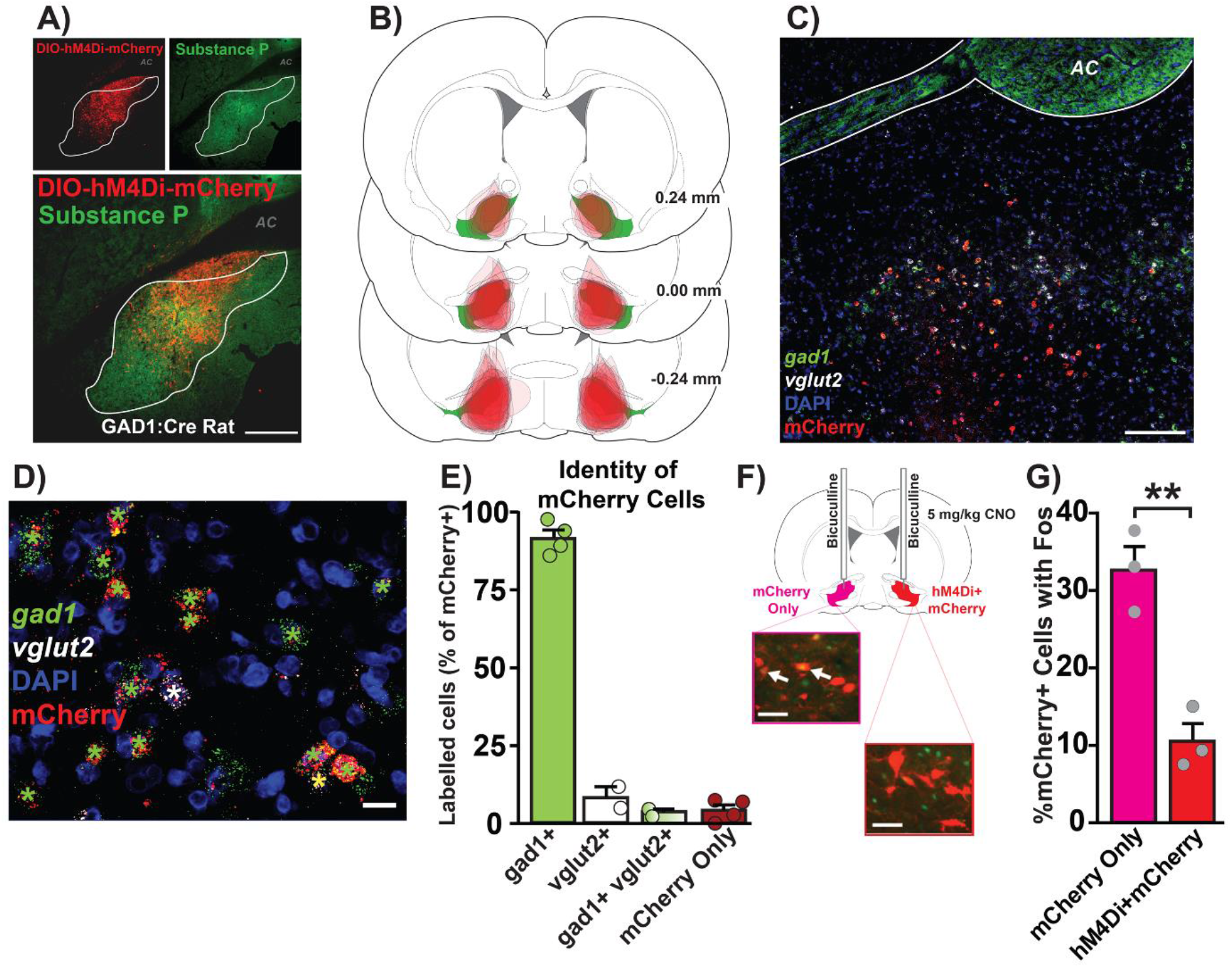
Anatomical, cellular, and functional characterization of hM4Di DREADDs in VP^GABA^ neurons. **A)** Localization of DIO-hM4Di-mCherry in substance P-defined VP borders. Top left panel: DIO-hM4Di-mCherry in VP. Top right panel: substance P demarcates VP from surrounding basal forebrain. Bottom: Merged DIO-hM4Di-mCherry and substance P image. AC = anterior commissure. Scale bar = 400 μm. **B)** Mapping of viral expression for each individual rat expressing hM4Di DREADDs. Numbers represent rostral/caudal coordinates relative to bregma. Green = substance P-defined VP. Red=DIO-hM4Di-mCherry expression. **C)** RNAscope fluorescent in situ hybridization for gad1, vglut2 and mcherry mRNA, with DAPI co-stain. Scale bar = 200 μm. **D)** Higher magnification of mRNA signal; Scale bar = 20 μm. Green star: mCherry+gad1; white star: mCherry+vglut2; yellow star: mCherry+gad1+vglut2. **E)** Identity of mCherry cells in VP. mCherry co-localized largely with gad1 mRNA (green bar), with few mCherry+ neurons expressing vglut2 mRNA. A small population of mCherry+ neurons expressed both gad1 and vglut2 (green+white gradient), and some cells lacked observable gad1 or vglut2 mRNA and only expressed mCherry (red). **F)** Schematic illustrating bilateral bicuculline (0.01 μg/0.5 μL) microinjection and systemic CNO (5 mg/kg) in rats with ipsilateral mCherry and contralateral hM4Di+mCherry in VP^GABA^ neurons (left image, Fos = green, red = mCherry; right image, Fos = green, red = hM4Di+mCherry). White arrows indicate colocalization of Fos in mCherry+ neurons. Scale bars = 40 μm. **G)** CNO reduced %mCherry+ cells colocalized with Fos in hM4Di+mCherry neurons, compared with contralateral mCherry only neurons. \*\**p* < 0.01, independent sample *t*-test. Each graph depicts mean + SEM, with dots representing individual rats.

To verify that hM4Di DREADDs measurably inhibit neural activity in VP^GABA^ neurons, we administered CNO systemically to GAD1:Cre rats (*n* = 3) with unilateral VP GAD1-dependent expression of hM4Di+mCherry, and contralateral VP GAD1-dependent mCherry only. We then pharmacologically disinhibited VP neurons bilaterally, using microinjections of the GABA_A_ antagonist, bicuculine (0.01 μg/0.5 μL), which robustly induces VP Fos (Smith and Berridge, 2005; Turner et al., 2008). As expected, fewer mCherry+Fos VP neurons were found in the hM4Di-expressing hemisphere than the mCherry hemisphere (**Fig 1F**; *t*_2_ = 18.12, *p* = 0.003), despite the fact that cannulae localizations were equivalent in each hemisphere. These results demonstrate that CNO, via actions at hM4Di, is capable of suppressing Fos in pharmacologically disinhibited VP^GABA^ cells, presumably by recruiting endogenous G_i/o_ signaling (Pleil et al., 2015; Roth, 2016).

### Inhibiting VP^GABA^ Neurons Reduces Risky Choices

Rats (*n* = 45) performed the risk task as expected, shifting their choices from the large reward when chance of shock was low, to the smaller but unpunished reward as the probability of shock increased (**Fig 2A**; GAD1:Cre rats main effect of block: *F*_(4, 96)_ = 40.68, *p* < 0.0001; rats: *F*_(4, 32)_ = 13.4, *p* < 0.0001). Male rats required higher average shock intensities than female rats (**Fig 2B**, *t*_40_ = 5.6, *p* < 0.0001), as reported previously (Orsini et al., 2016). However, after this individualized shock titration, males and females performed equivalently on the task; similarly shifting their choice from the large/risky to the small/safe reward option as shock probability increased (block: *F*_(4, 179)_ = 76.5, *p* < 0.0001). Importantly, no sex differences were detected for percent choice of the risky option (**Fig 2C**, sex: *F*_(1, 179)_ = 0.19, n.s.; block x sex interaction: *F*_(4, 179)_ = 0.07, n.s.). Shock intensity required for stable behavior in each GAD1:Cre rat did not predict the subsequent magnitude of VP^GABA^ neuron inhibition effects; i.e. shock intensity did not correlate with CNO effects (CNO day -vehicle day) on large/risky lever pressing (*r* = 0.14, n.s.) nor on small/safe pressing (*r* = -0.05, n.s.).

**Figure 2.**
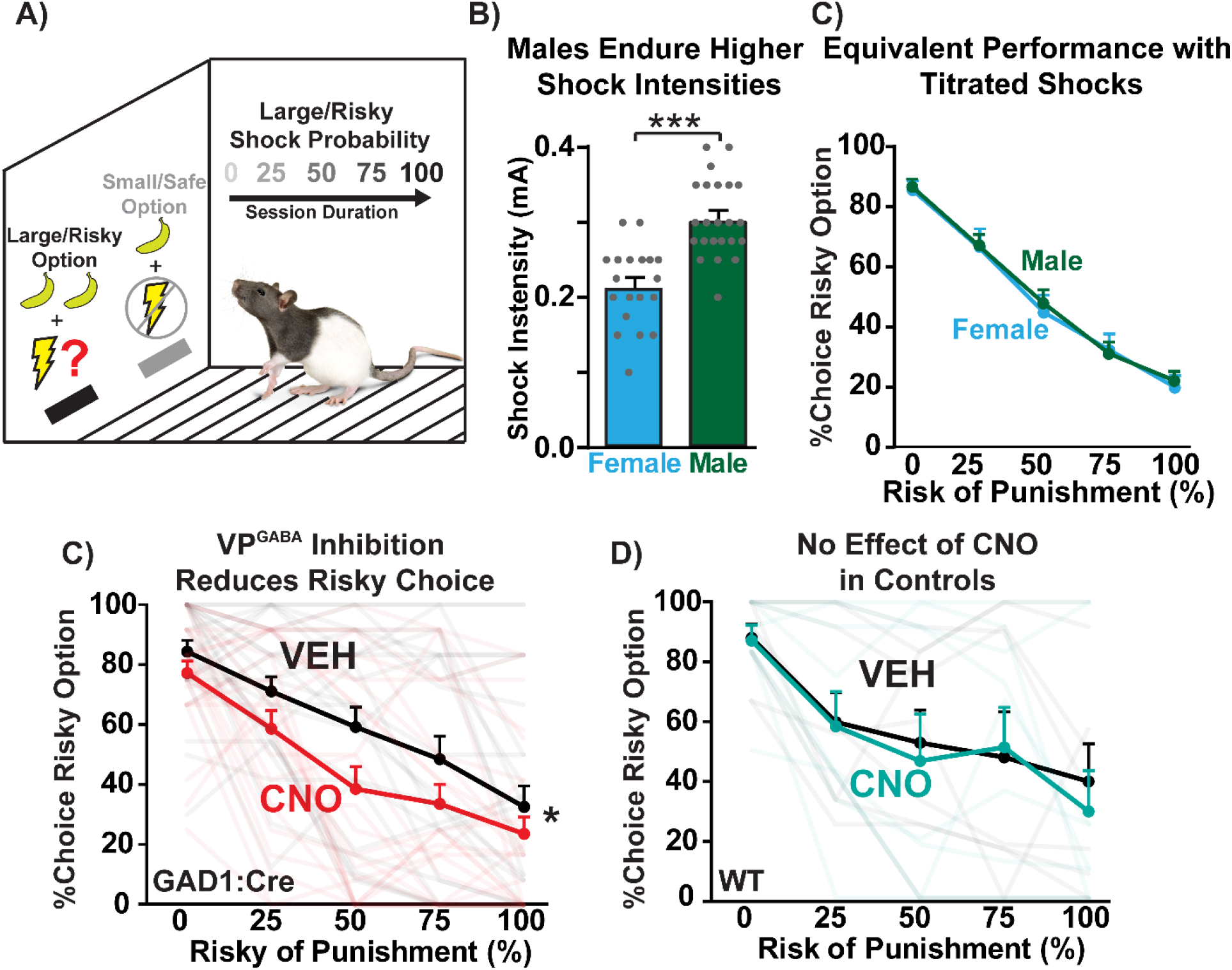
Inhibiting VP^GABA^ neurons reduces risky choice. **A)** Schematic of risky decision task modified from (Simon et al., 2009). Sessions consisted of forced choice trials (1 available option; small/safe or large/risky) and free choice trials (2 available options; small/safe and large/risky), with ascending footshock probability associated with selection of the large/risky reward option. **B)** Male rats required a higher shock intensity than females for appropriate performance of the risky decision task, as previously reported (Orsini et al., 2016). **C)** Equivalent average performance of male and female rats on the risky decision task with shock titration. **D)** GAD1:Cre rats administered CNO (red line) exhibit a decrease in %choice of the risky option relative to vehicle-treated rats (black line). **E)** No effect of CNO (teal line) in WT rats compared with vehicle treatment (black line). \*\*\**p* < 0.0001, Independent sample *t*-test; \**p* < 0.05, treatment main effect. Semi-transparent lines represent data from individual rats tested with CNO (red/teal) or vehicle (black). Each graph depicts mean + SEM.

In GAD1:Cre rats (*n* = 30), CNO reduced choice of the large, risky reward option (**Fig 2D**, treatment: *F*_(1, 24)_ = 4.62, *p* = 0.042). Though no significant overall treatment x block interaction was detected, block-specific comparisons revealed that suppression of choice of the large reward was statistically different during the 50% (Sidak: p = 0.0014) and 75% (Sidak: p = 0.038) blocks, but not at the 0, 25, or 100% shock probability blocks (all n.s.). To further probe effects of VP^GABA^ inhibition on the ability of positive or negative experiences to adjust ongoing behavior, we performed a win-stay/lose-shift analysis (Onge et al., 2011). CNO, relative to vehicle, did not alter the proportion of trials in which either win-stay (Vehicle: m ± SEM = 0.70 ± 0.055; CNO: m = 0.69 ± 0.036; *t*_18_ = 0.18, n.s.) or lose-shift occurred (Vehicle: m = 0.42 ± 0.062; CNO: m = 0.49 ± 0.076; *t*_18_ = 0.71, n.s.), indicating that the effects of VP^GABA^ inhibition on choice were not driven by altered sensitivity to outcomes of recent choices. As expected, latency to press the large/risky reward option increased as the footshock probability increased (one-way ANOVA for vehicle day data: *F*_(4, 118)_ = 7.21, p < 0.0001), which did not occur for latency to press the small/safe reward option (*F*_(4, 119)_ = 1.44 n.s.). CNO selectively increased the latency to press the large/risky reward option (**Fig 3A**, treatment: *F*_(1, 231)_ = 13.5, *p* = 0.0003), especially when the uncertainty of footshock was maximum; during the 50% footshock block (Sidak posthoc: *p* = 0.0053). In contrast, CNO failed to impact latency to press the small/safe reward option (**Fig 3B**, *F*_(1, 237)_ = 1.29, n.s.). CNO also increased the total number of omitted trials, especially in the blocks with the highest probability of shock (**Fig 3C**, treatment x block interaction: *F*_(4, 96)_ = 11.91, *p* < 0.0001; Sidak posthoc: 50% block: *p* = 0.0006; 75%: *p* < 0.0001; 100%: *p* < 0.0001). In addition, due to decreased pressing of the large/risky option and increased omissions, CNO-treated GAD1:Cre rats obtained fewer rewards overall than on vehicle day (treatment: *F*_(1, 24)_ = 6.95, *p* = 0.015, data not shown).

**Figure 3.**
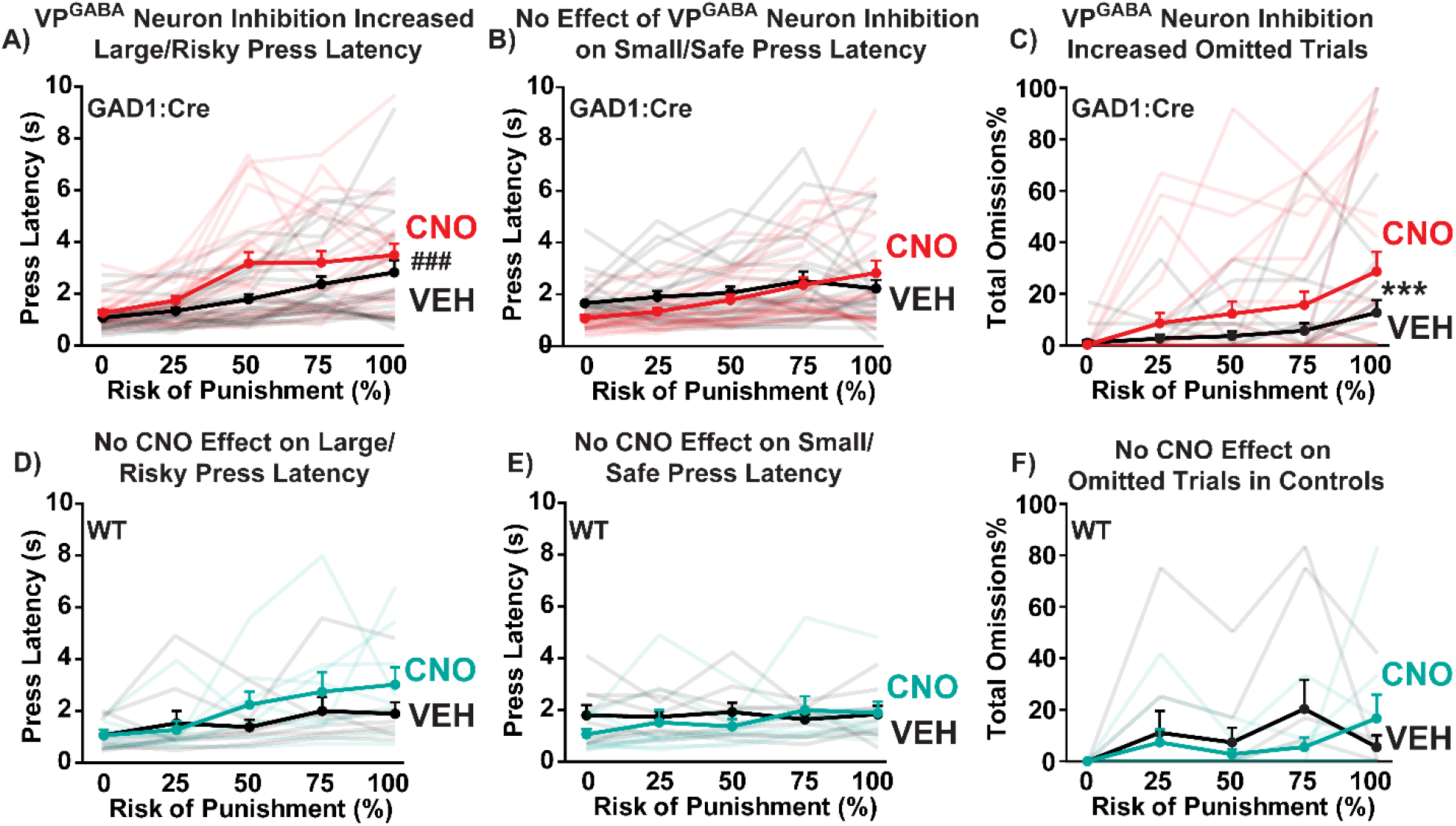
VP^GABA^ neuron inhibition increases latency to select the large/risky option, and trial omissions. **A-C)** In GAD1:Cre rats, CNO (red lines) increased the latency to press the large/risky reward lever, relative to vehicle day in the same rats (black lines; ###*p* < 0.001, treatment main effect). **B)** In contrast, CNO in GAD1:Cre rats did not affect latency to press the small/safe reward lever. **C)** CNO in GAD1:Cre rats increased the percentage of trials omitted on high-risk blocks (\*\*\**p* < 0.001, treatment x block interaction). **D-F)** In WT rats without VP DREADDs, CNO (teal lines) did not alter **D)** omissions, **E)** latency to press the large/risky reward lever, **F)** or latency to press the small/safe reward option, relative to vehicle day (black lines). Semi-transparent lines represent data from individual rats tested with CNO (red/teal) or vehicle (black). Each graph depicts mean + SEM.

### VP^GABA^ Inhibition Effects are Specific to Risky Choices: No Effect on Reward Magnitude Discrimination

#### Reward Magnitude Discrimination

A separate group of GAD1:Cre rats (*n* = 8) were trained as described above, but in the absence of shock, to confirm their ability to discriminate reward magnitude after VP^GABA^ neuron inhibition. As expected, rats nearly exclusively chose the large reward over the small one during training, and after vehicle treatment (vehicle day: large versus small reward: *t*_7_ = 11.11, p < 0.0001). After CNO, GAD1:Cre rats showed a nearly identical preference as on their vehicle test day (**Fig 4A,** treatment: *F*_(1, 56)_ = 1.08, n.s.), showing that VP^GABA^ inhibition does not affect rats’ preference for a large reward over a small one. This said, as in the task where the larger reward was associated with a probabilistic shock, CNO increased omitted trials (**Fig 4B,** treatment x block interaction: *F*_(4, 24)_ = 4.0, *p* = 0.013), and decreased total rewards obtained (**Fig 4C,** treatment x block interaction *F*_(4, 24)_ = 3.73, *p* = 0.017), consistent with an overall reduction in motivation. Posthoc tests revealed that CNO increased omissions only in the 3^rd^ block (Sidak posthoc: *p* < 0.01) and 4^th^ block (*p* < 0.05), and similarly only decreased rewards obtained in the 3^rd^ (Sidak posthoc: *p* < 0.01) and 4^th^ (*p* < 0.05) blocks. Emergence of satiety in later blocks likely accounts for why the 5^th^ block of trials converged for both omissions and rewards obtained, as a significant main effect of block was observed for both omissions (*F*_(4, 24)_ = 21.4, *p* < 0.001) and rewards obtained (*F*_(4, 24)_ = 19.4, *p* < 0.001). However, CNO did not impact choice latency relative to vehicle (treatment: *F*_(1, 59)_ = 0.86, n.s.), unlike in the shock version of the task where CNO selectively increased latency to press the large/risky lever. The lack of effect on choice latencies induced by VP^GABA^ neuron inhibition in this experiment suggest that the increased decision times observed on the risky decision task were not attributable to a generalized psychomotor slowing, but rather to increased deliberation time in weighing the costs and benefits of the risky choice.

**Figure 4.**
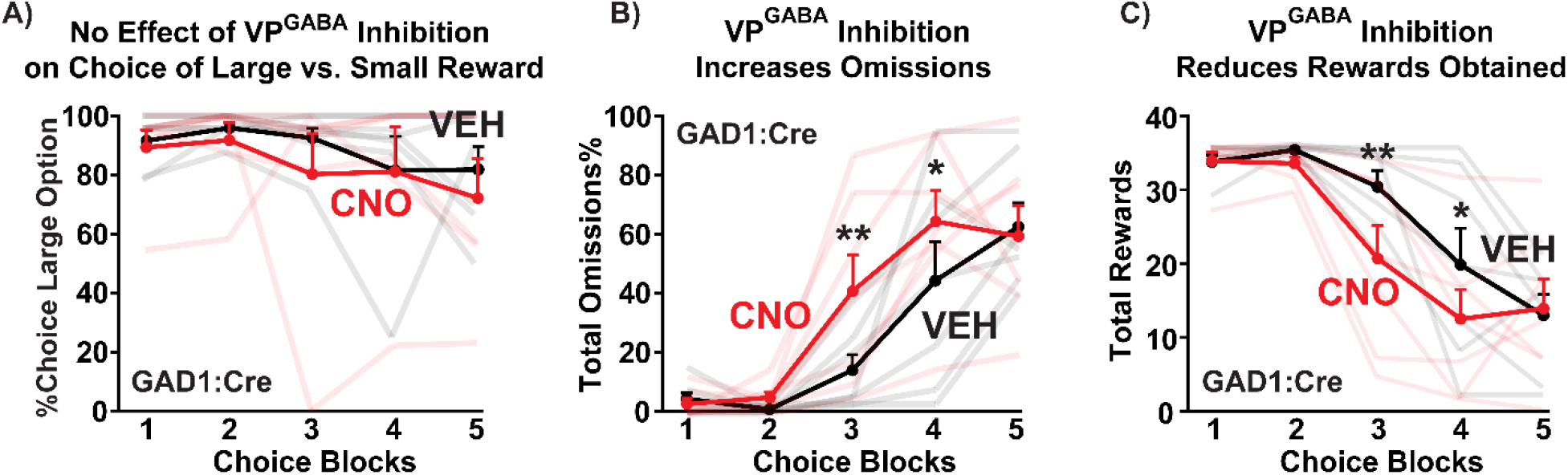
Inhibiting VP^GABA^ neurons spares the ability to choose between large and small rewards, while decreasing motivation. **A)** In the absence of shock punishment, CNO-treated GAD1:Cre rats (red line) showed no change in percentage choice of the large (2 pellets) versus small (1 pellet) reward option, compared with vehicle treatment (black line). **B)** GAD1:Cre rats omitted more trials during CNO tests (red line) compared with vehicle treatment (black line). **C)** CNO treatment in GAD1:Cre rats (red line) reduced rewards obtained relative to vehicle (black line). Semi-transparent lines represent data from individual rats tested with CNO (red/teal) or vehicle (black). Each graph depicts mean + SEM. \**p* < 0.05, \*\**p* < 0.01, Sidak posthoc tests.

### Inhibiting VP^GABA^ Neurons Suppresses Instrumental Responding for High Value Foods, without Impairing General Locomotion

#### Similar Suppression of Palatable Food Responding During Hunger and Satiety

We examined effects of inhibiting VP^GABA^ neurons on low effort (FR1) operant responding for highly palatable banana pellets. When GAD1:Cre rats (*n* = 10) were tested under mild food restriction, CNO reduced active port responding (**Fig 5A**: *t*_9_ = 2.58, *p* = 0.03), without affecting inactive port responding (*t*_9_= 0.79, n.s.). Effects of VP^GABA^ neuron inhibition were similar when rats were tested in the same manner while maintained on *ad libitum* chow 23 hrs/day (**Fig 5B**, active port responses: *t*_9_ = 2.65, *p* = 0.027, inactive: *t*_9_ = 0.97, n.s.), indicating that VP^GABA^ neurons are required for low effort instrumental pursuit of a highly salient, palatable reward regardless of physiological need state.

**Figure 5.**
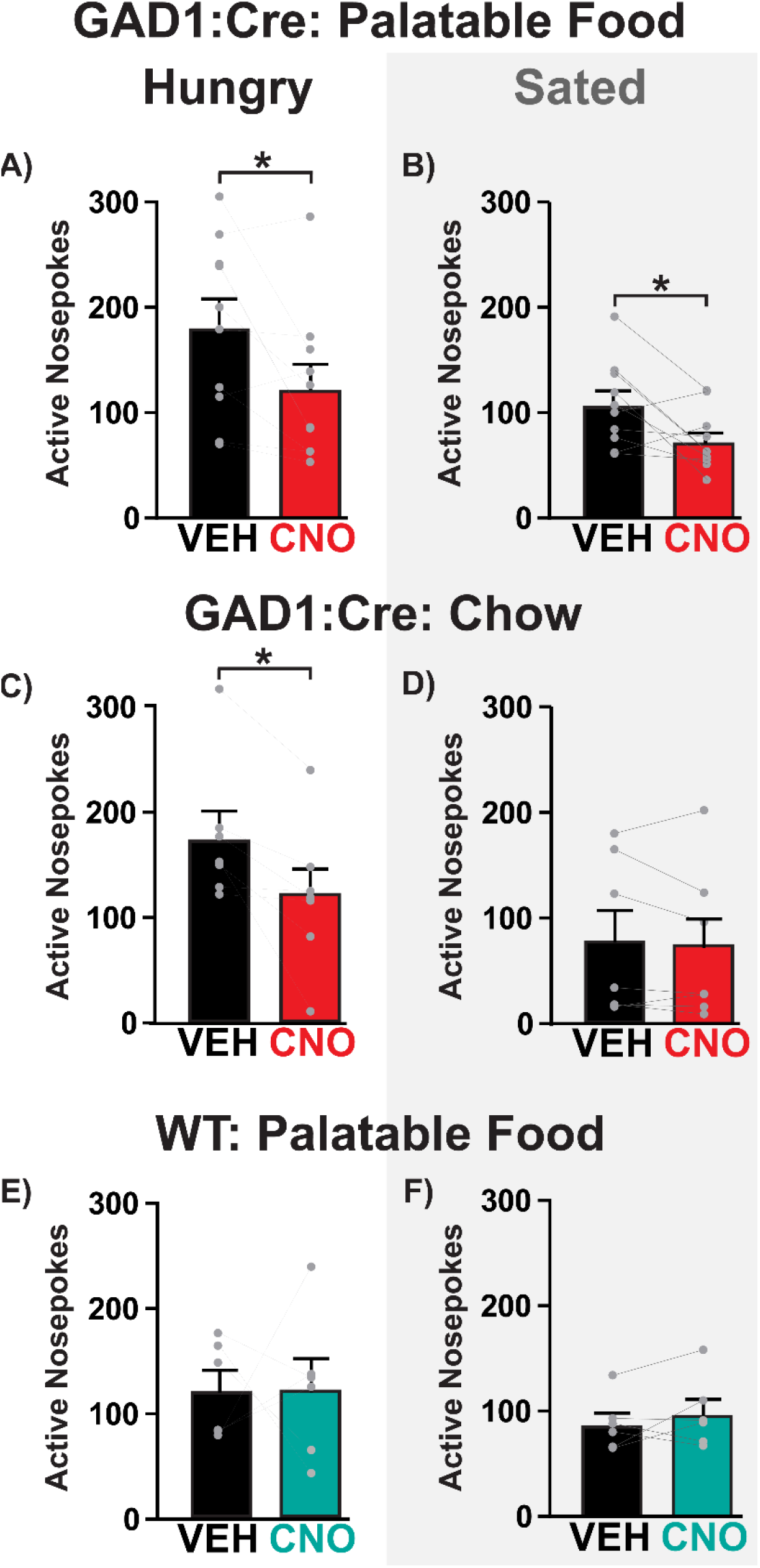
Inhibiting VP^GABA^ neurons reduces responding for palatable food and chow in hungry rats, but only reduces responding for palatable food (not chow) in sated rats. **A-B)** In GAD1:Cre rats, CNO (red bars) reduced FR1 active nosepokes for palatable food in **A**) hungry and **B**) sated rats, relative to vehicle treatments (black bars). **C-D)** CNO in GAD1:Cre rats (red bars) reduced FR1 active nosepokes for chow relative to vehicle (black bars) in **C**) hungry, but not **D**) sated rats. **E-F**) No effect of CNO on active nosepokes in WT controls (teal bars) relative to vehicle treatment (black bars) under **E**) hungry or **F**) sated conditions. \**p* < 0.05, paired sample *t*-test. Each graph depicts mean + SEM, and dots represent individual rats.

#### Suppression of Responding for Less-Palatable Chow Only During Hunger

We next examined effects of inhibiting VP^GABA^ neurons on low effort (FR1) operant responding for standard chow pellets under hunger and satiety conditions (*n* = 8). Inhibiting VP^GABA^ neurons reduced active port responding for chow when rats were hungry (**Fig 5C**, *t*_6_ = 3.12, *p* = 0.021), but not when they were fed ad libitum (**Fig 5D**, *t*_6_ = 0.89, n.s.), as shown by the significant interaction between hunger state and vehicle/CNO treatment (*F*_1, 6_ = 6.31, *p* = 0.046). However, we note that responding for chow during satiety was quite low in some animals, raising the possibility of a floor effect.

#### Robust Suppression of High-Effort Palatable Food Seeking

When GAD1:Cre rats (*n* = 22) were trained on a progressive ratio to stably respond for palatable banana pellets, CNO suppressed breakpoint (**Fig 6A-B**, *t*_21_ = 2.4, *p* = 0.026), and trended toward suppressing active port responses (vehicle: m = 1050 ± 136.7, CNO: m = 847.5 ± 122.4; *t*_21_ = 2.0, *p* = 0.059). The low number of inactive port responses was unaffected (vehicle: m = 15.8 ± 2.9, CNO: m = 18.3 ± 4.0; *t*_21_ = 0.53, n.s.).

**Figure 6.**
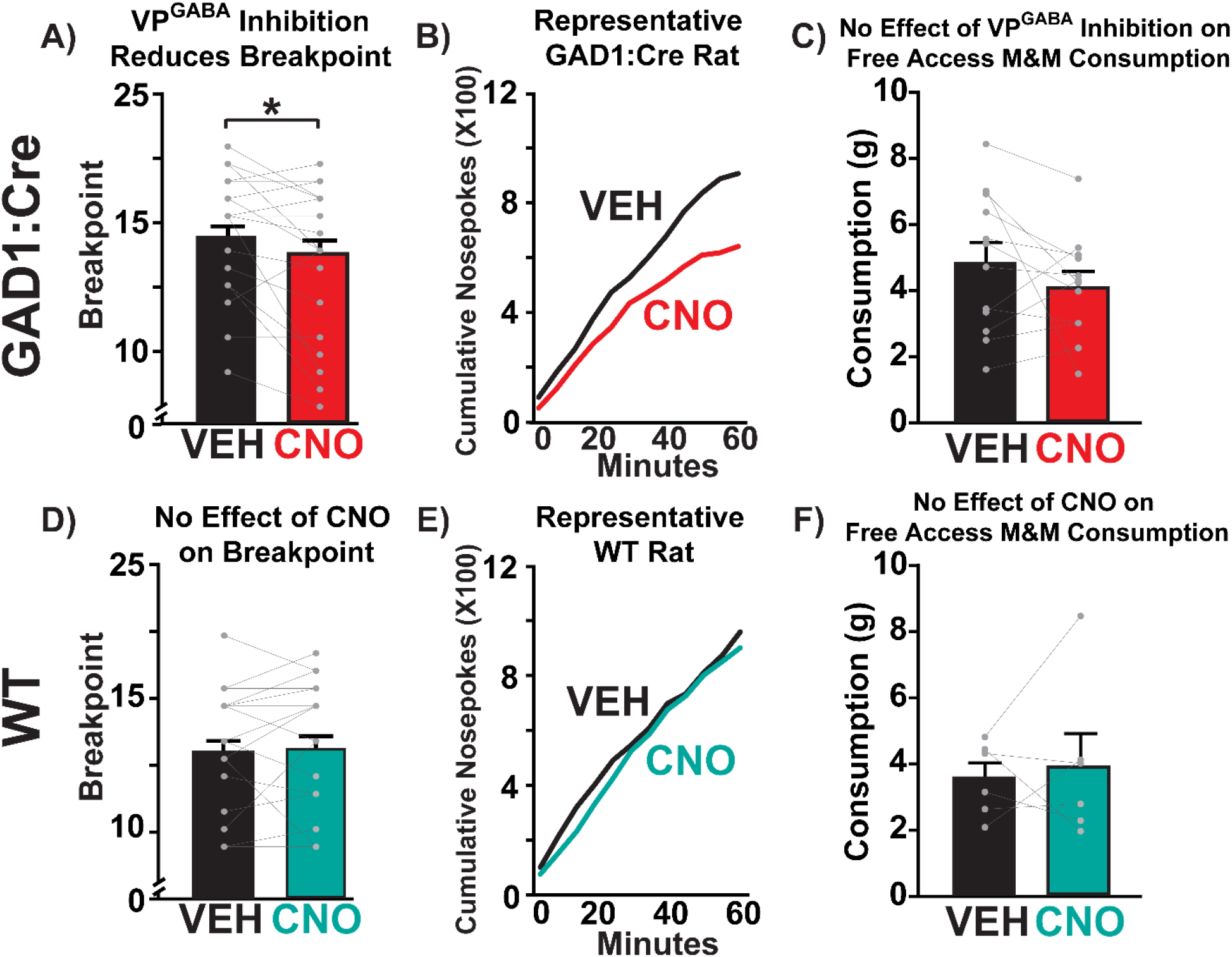
Inhibiting VP^GABA^ neurons reduces progressive ratio motivation for palatable food, without impairing free access palatable chocolate intake. **A)** In GAD1:Cre rats, CNO (red bar) reduces breakpoint relative to vehicle (black bar). **B)** Cumulative nosepokes for a representative GAD1:Cre rat during vehicle (black line) and CNO (red line) progressive ratio tests are shown. **C)** CNO in GAD1:Cre rats (red bar) fails to alter 1 hr free access M&M consumption, relative to vehicle test (black bar). **D)** In WT controls, CNO (teal bar) does not affect breakpoint compared with vehicle day (black bar). **E)** Cumulative nosepokes for representative WT rat during vehicle (black line) and CNO (teal line) progressive ratio tests are shown. **F)** CNO in WT rats (teal bar) fails to alter 1 hr free access M&M consumption, relative to vehicle test (black bar). \**p* < 0.05, paired sample *t*-test. Each graph depicts mean + SEM, and dots represent individual rats.

#### Non-Operant Spontaneous Intake of Palatable Food is Unaffected

To determine effects of VP^GABA^ neuron inhibition on spontaneous intake of a highly palatable sweet and fatty food meal (*n* = 12), we examined 2 hr intake of peanut butter M&M™ candies, placed directly on the floor of a familiar testing chamber. CNO failed to affect intake (g) in GAD1:Cre rats (**Fig 6C**, *t*_11_ = 1.24, n.s.).

#### Locomotor Activity

Effects of VP^GABA^ inhibition with CNO treatment failed to alter either horizontal locomotion or rearing behavior in GAD1:Cre rats (*n* = 12) (Distance travelled: vehicle m = 12026 ± 1006 cm, CNO m = 13185 ± 1601 cm, *t*_11_ = 0.71, n.s.; Rearing: vehicle m = 183.6 ± 17.1 rears, CNO m = 188 ± 22.7 rears, *t*_11_ = 0.20, n.s.).

### Inhibiting VP^GABA^ Neurons Decreases Motivation to Avoid Footshock Without Impacting Motor or Affective Reactions to Shock

#### Latency to Avoid Footshock Increases after VP^GABA^ Neuron Inhibition

VP^GABA^ neuron inhibition suppressed operant risky decision making and food seeking, so we next sought to determine whether this manipulation also affects negatively reinforced operant responding (*n* = 11). CNO did not affect the overall propensity of rats to avoid shocks rather than to escape them (**Fig 7A**, change from baseline avoidance%: *t*_10_ = 1.50, n.s.; Vehicle: m = 38.82 ± 5.71; CNO: m = 28.96 ± 5.07; raw %avoidance: *t*_10_ = 2.2, *p* = 0.053), suggesting that their general strategy was not altered by this manipulation. However, CNO selectively increased latency to press to avoid shock in GAD1:Cre rats (**Fig 7B**, *t*_10_ = 2.60, *p* = 0.027), consistent with reduced motivation to avoid the impending, signaled shock. Escape latency was not similarly impacted by CNO (**Fig 7C**, *t*_10_ = 1.36, n.s.), indicating that rats were still fully capable of pressing to terminate an ongoing shock.

**Figure 7.**
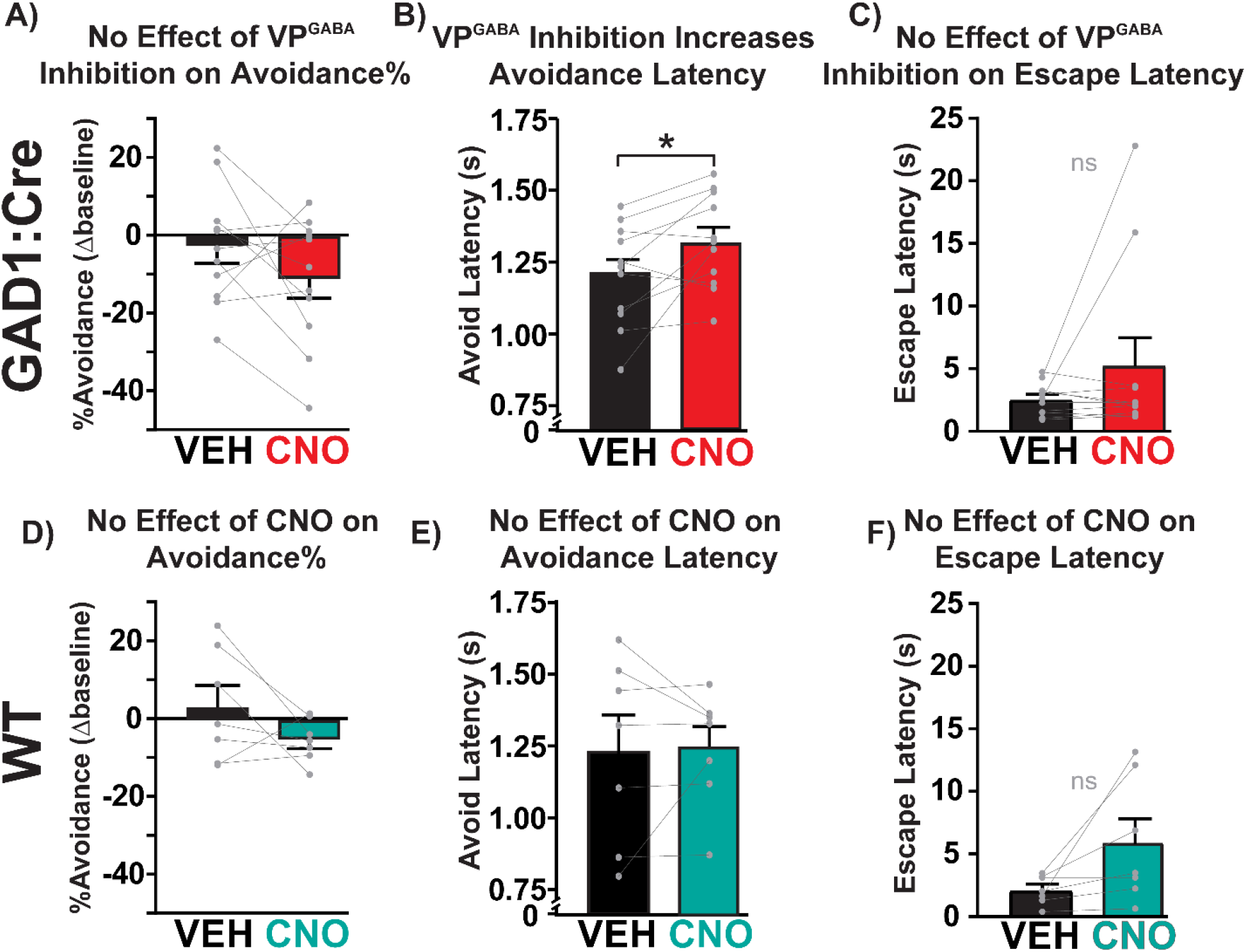
Inhibiting VP^GABA^ neurons increases avoidance latency, without affecting avoidance propensity or escape latency. **A)** CNO did not affect avoidance% in GAD1:Cre rats (red bar) relative to vehicle treatment (black bar) (change in avoidance% on vehicle/CNO test from the day preceding each treatment). **B)** CNO increased latency to lever press to avoid being shocked in GAD1:Cre rats (red bar) compared with vehicle treatment (black bar). **C)** No effect of CNO on escape latency in GAD1:Cre rats. **D-F)** CNO in WT controls (teal bars) failed to impact **D)** avoidance%, **E)** avoidance latency, or **F)** escape latency relative to vehicle treatments (black bars). \**p* < 0.05, paired sample *t*-test. Each graph depicts mean + SEM, and dots represent individual rats.

#### No Effects on Motor or Affective Responses to Shock

As expected, shock-induced motor reactivity scores parametrically increased with footshock intensity (**Fig 8A,** GAD1:Cre block: *F*_(6, 112)_ = 100.9, *p* < 0.0001). Motor reactivity scores were not affected by CNO in GAD1:Cre rats (*n* = 11) (treatment: *F*_(1, 112)_ = 0.27, n.s.), nor was the maximum shock intensity endured altered (vehicle m = 0.27 ± 0.017 mA; CNO m = 0.29 ± 0.014 mA, *t*_10_ = 0.94, n.s.).

**Figure 8.**
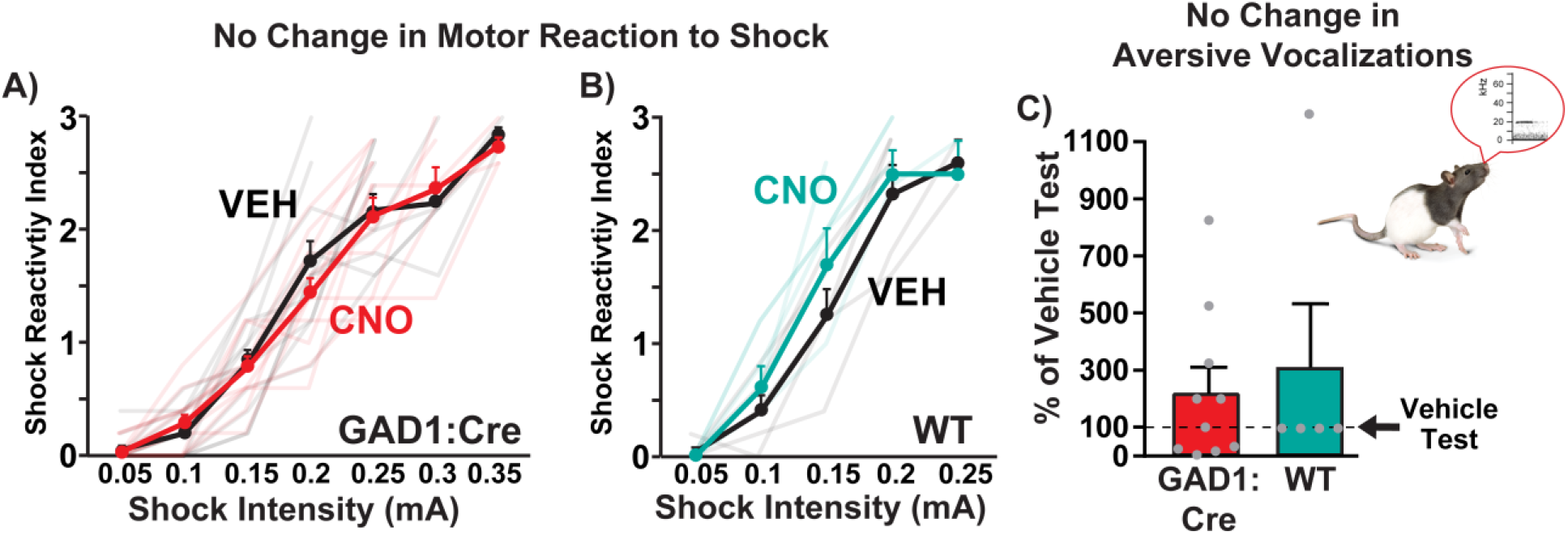
Motor and affective reactions to footshock unaffected by inhibiting VP^GABA^ neurons. **A)** CNO in GAD1:Cre (red line) or in **B)** WT rats (teal line) did not impact shock reactivity index associated with ascending intensity footshocks, relative to vehicle rats (black lines). **C)** CNO in GAD1:Cre rats (red bar) or WT controls (teal bar) failed to alter aversion-related ultrasonic vocalizations during high intensity footshock (0.75 mA) sessions (percent of vehicle test day). Each graph depicts mean + SEM, and dots represent individual rats.

#### No Effect on Shock-Induced Negative Affective Vocalizations

We also examined aversion-related 22 kHz ultrasonic vocalizations emitted in response to repeated, moderate intensity shocks (0.75 mA/1 sec, delivered every min for 5 min). These USVs were in the frequency range of well-characterized aversion-related USVs (Knutson et al., 2002; Portfors, 2007; Mahler et al., 2013b), with a mean frequency of m = 23.5 ± 1.1 kHz, and a mean duration of m = 1306.4 ± 142.1 ms. CNO failed to alter the number of 22 kHz vocalizations emitted in GAD1:Cre rats (*n* = 10) (**Fig 8C**, % of vehicle day USVs, compared to 100% with one sample *t*-test: *t*_9_ = 1.5, n.s.). We also observed some vocalizations >30 kHz, linked to positive affect (Knutson et al., 2002; Portfors, 2007; Brudzynski, 2013). These vocalizations, however, occurred largely during the 2 min pre-footshock baseline (high frequency USVs/min on vehicle test for GAD1:Cre and WT: m = 46.4 ± 14.8) compared to the subsequent 5 min intermittent footshock period (m = 9.6 ± 5.8; pre-footshock vs. footshock period: *t*_14_=3.16, *p* = 0.0069). Production of these high frequency vocalizations was also unaffected by CNO treatment (% of vehicle day USVs, compared to 100% with one sample *t*-test: *t*_9_ = 1.74, n.s.).

### Minimal DREADD-Independent Effects of CNO

Across all nine behavioral tasks implemented here, we saw few non-specific effects of CNO in WT rats lacking DREADD expression. In the risky decision task, administering CNO to WT rats did not affect risky choice (**Fig 2E**, treatment x block ANOVA, no main effect of treatment: *F*_(1, 8)_ = 0.055, n.s.), though an overall ANOVA did not reveal a significant three-way interaction of treatment, genotype, and block (*F*_(4, 154)_ = 0.47, n.s.), or a genotype x treatment interaction (*F*_(1,32)_ = 0.61, n.s.), likely due to the relatively low WT rat group size. In WT rats, CNO had no effect on total omissions (**Fig 3F**, treatment: *F*_(1, 8)_ = 0.60, n.s.), total rewards obtained (treatment: *F*_(1, 8)_ = 0.044, n.s.), or latency to press for either large/risky (**Fig 3D**, treatment: *F*_(1, 77)_ = 2.68, n.s.) or small/safe rewards (**Fig 3E,** treatment: *F*_(1, 77)_ = 0.91, n.s.), though no significant genotype x treatment x block interactions were detected for these variables (*p* > 0.05). CNO also failed to alter FR1 responding for palatable pellets in either food-deprived (**Fig 5E**, active vehicle versus CNO: *t*_5_ = 0.036, n.s.; inactive: *t*_5_ = 1.04, n.s.) or sated WT rats (**Fig 5F**; active: *t*_5_ = 1.01, n.s.; inactive: *t*_5_ = 1.04, n.s.), and a significant genotype x treatment interaction further demonstrate this DREADD-specific effect in sated rats (genotype x treatment interaction; active lever: *F*_(1, 14)_ = 5.70, *p* = 0.032), though not significantly so for food-deprived ones (genotype x treatment interaction; active lever: *F*_(1, 14)_ = 1.89, n.s.). Progressive ratio responding for palatable pellets was also unaffected by CNO in WT rats (**Fig 6D-E**, breakpoint: *t*_16_ = 0.63, n.s.; active nosepokes: *t*_16_ = 0.54, n.s.; inactive nosepokes: *t*_16_= 0.59, n.s.), as was spontaneous M&M consumption (**Fig 6F**, *t*_5_ = 0.41, n.s.). A significant genotype x treatment interaction was found for breakpoint on the progressive ratio task (genotype x treatment interaction breakpoint: *F*_(1, 37)_ = 4.49, *p* = 0.041), but this was not significant for spontaneous M&M consumption (genotype x treatment interaction: *F*_(1, 16)_ = 1.11, n.s.). Likewise, CNO in WT rats did not impact general locomotion (vehicle m = 13621 ± 2580 cm, CNO m = 16768 ± 3039 cm, *t*_5_ = 1.02, n.s.) or rearing (vehicle m = 197.8 ± 29.7 rears, CNO m = 174 ± 26.7 rears, *t*_5_ = 0.56, n.s.), and the genotype x treatment interactions were also non-significant (locomotion: *F*_(1, 16)_ = 0.40, n.s.; rearing: *F*_(1, 16)_ = 0.42, n.s.).

Shock-related behaviors were also largely unaffected in WTs by CNO, including motor reactions to shock (**Fig 8B**, treatment: *F*_(1, 33)_ = 0.85, n.s.), maximum shock intensity tolerated (vehicle: m = 0.22 ± 0.012; CNO: m = 0.21 ± 0.019; *t*_4_ = 1.0, n.s.), avoidance propensity (**Fig 7D**, change from baseline avoidance%: vehicle day raw percentage: m = 42.08 ± 10.73, CNO day raw percentage: m = 28.96 ± 9.07, *t*_6_=1.54, n.s.), or avoidance latency (**Fig 7E**, *t*_6_ = 0.064, n.s.), though raw avoidance% was modestly decreased by CNO in WT rats (*t*_6_ = 2.9, *p* = 0.027). No significant genotype x treatment interaction was found for either avoidance propensity (*F*_(1, 16)_ = 0.0002, n.s.) or avoidance latency (*F*_(1, 16)_ = 1.79, n.s.). Both 22 kHz (aversion-related) and >30 kHz (positive affect-related) USVs were unchanged by CNO in WT rats (**Fig 8C**, % of vehicle day USVs, compared to 100% with one sample *t*-test, 22kHz: *t*_4_ = 1.0, n.s.; >30 kHz: *t*_4_ = 0.96, n.s.). CNO in WT rats trended towards increasing latency to escape (**Fig 7F**, *t*_6_ = 2.2, *p* = 0.07), though no genotype x treatment interaction was found (*F*_(1, 16)_ = 0.14, n.s.).

## Discussion

Here we show that VP^GABA^ neurons play a fundamental role in high-stakes motivation, and thereby affect risky decision making strategies. Engaging G_i/o_ signaling in VP^GABA^ neurons with DREADDs interfered with both operant pursuit of desirable foods, as well as operant response to cancel an impending shock. In contrast, VP^GABA^ neurons play no apparent role in pursuit of less valuable food, in spontaneous food consumption, or in affective responses to shock itself. This selective VP^GABA^ neuron involvement in motivated operant responding may therefore extend beyond the pursuit of rewards, into avoidance of harm. Accordingly, when both opportunity and risk are present (as is usually the case in the natural world), VP^GABA^ inhibition biased decision making toward a more conservative, risk-averse strategy. Collectively, these results show that VP^GABA^ neurons crucially influence high-stakes decision making, and thus likely contribute to both the normal desires of life, and to darker pursuits in those with disorders of impaired judgement like addiction.

### VP^GABA^ Neuron Inhibition Promotes Conservative Decision Making by Suppressing Motivation

In a risky decision making task, chemogenetically inhibiting VP^GABA^ neurons promoted selection of a small but safe option over a large but risky one, without impairing the ability to discriminate between rewards of different magnitudes. VP^GABA^ inhibition also increased trial omissions and decreased the number of rewards obtained in the presence or absence of shock—consistent with decreased motivation for food. Similar increases in latency and omissions have been shown following optogenetic inhibition of all VP neurons in operant assays of sucrose seeking (Richard et al., 2016). Yet VP^GABA^ inhibition effects were not merely motivational in nature—food seeking was not indiscriminately suppressed. Instead, VP^GABA^ inhibited rats shifted more readily to a small but safe reward option, avoiding the large but risky one, even when the risk of shock was relatively low. Moreover, when rats did select the large/risky choice, VP^GABA^ inhibition caused them to deliberate longer—an effect which was not present on trials when the small/safe option was chosen. In contrast, when VP^GABA^ neuron inhibition occurred in a low stakes (no shock) version of the task, no such effects on choice latency were seen. In other words, inhibiting VP^GABA^ neurons seemed to selectively promote a more conservative, risk-averse decision making strategy by suppressing appetitive motivation.

Of course, VP does not act alone to influence risky choice, but rather within wider mesocorticolimbic circuits that integrate motivational states with encountered opportunities and threats, in pursuit of generating maximally adaptive behavior under motivational conflict. Indeed, numerous brain regions contribute to risky decision making in rats, including prefrontal cortices, basolateral amygdala, lateral habenula, ventral tegmental area, and nucleus accumbens (NAc) (Floresco et al., 2008; Orsini et al., 2015b). Notably, lateral orbitofrontal cortex lesions have similar effects on latency and propensity to make risky choices as VP^GABA^ neuron inhibition did here (Orsini et al., 2015a), implying functional, if not direct anatomical interactions between these structures (Simmons et al., 2014). Interestingly, activating D2 dopamine receptors in VP’s largest afferent input, the GABAergic NAc, similarly promotes risk-averse behavior in adolescent rats (Mitchell et al., 2014). Though infusion of a D2 agonist in NAc would likely disinhibit (excite) VP neurons (Gallo et al., 2018), paradoxically we find here that *inhibiting* VP^GABA^ neurons with DREADDs causes a similarly risk-averse phenotype. Reconciling these findings is an important future direction, and could involve experience-related plasticity in D1 (i.e. “direct pathway”) versus D2 (i.e. “indirect pathway”) inputs from NAc (Kupchik et al., 2015; Creed et al., 2016; Heinsbroek et al., 2017; O’Neal et al., 2019), differences between adolescent and adult decision making processes (Spear, 2000), currently-unknown specificity of NAc inputs to VP cell subpopulations (e.g. glutamate versus GABA), or potentially non-NAc inputs to VP that may influence reward-seeking decisions (Richard et al., 2016; Ottenheimer et al., 2018).

### Role for VP^GABA^ Neurons in Seeking High Value Food, without Affecting Food Consumption

Having found that VP^GABA^ neuron perturbation stifled risky choice, we next sought to determine how inhibiting these neurons impacts “pure” tests of food seeking and intake, in the absence of potential harm. VP’s role in food ingestion and hedonics has been known for decades (Morgane, 1961; Stratford et al., 1999; Castro et al., 2015), though how VP neuronal subtypes participate in this was unclear. Here, we show that chemogenetically inhibiting VP^GABA^ neurons suppresses operant pursuit of high-value foods like palatable pellets under both low and high effort conditions. In contrast, pursuit of less palatable chow was affected by VP^GABA^ inhibition only when this food was valued because rats were hungry. These results suggest that VP^GABA^ neurons selectively promote seeking of high-value rewards, regardless of whether value is instantiated by the inherent palatability of the food, by the presence of hunger, or by the necessity to pay a cost such as effortful responding, or potential for shock.

Interestingly, whereas inhibiting VP^GABA^ neurons decreased operant pursuit of valuable food rewards, it did not impair spontaneous consumption of palatable chocolate, suggesting that these neurons mediate instrumental *seeking* of high value rewards, but not necessarily *consumption* of the reward, once obtained. Neurobiological dissociation between seeking and consumption has been previously shown within ventral striatal networks (Berridge and Robinson, 2003). For example, intra-NAc dopamine antagonism diminishes operant reward seeking, but leaves reward consumption unimpaired (Kelley et al., 2005; Salamone and Correa, 2012). Similarly, inhibiting VP impairs conditioned food or salt seeking, without impacting unconditioned consumption of these rewards (Farrar et al., 2008; Chang et al., 2017). Our results extend these findings, showing that VP^GABA^ neurons in particular are required for pursuit, but not consumption of food. This said, VP stimulation with opioid agonist or GABA antagonist drugs robustly increases chow consumption, and opioid drugs also enhance hedonic reactivity to sweet tastes (Smith and Berridge, 2005, 2007). In addition, VP lesions suppress all food intake, and lesioned animals will starve without forced feeding (Cromwell and Berridge, 1993). Given these findings, the present results could suggest lack of VP^GABA^ neuron involvement in these consummatory effects, or they could be a product of mechanistic differences between DREADDs and lesions, or other unknown factors.

### VP^GABA^ Neurons and Appetitive versus Aversive Motivation

A recent surge of studies suggest that VP^GABA^ neurons promote appetitive behavior and reward, whereas intermingled VP glutamate neurons instead mediate behavioral withdrawal and aversion. For example, VP^GABA^ neurons fire in response to water rewards and their predictors in mice, especially when those rewards are particularly valuable due to thirst (Stephenson-Jones et al., 2020). Optogenetic activation of mouse VP^GABA^ neurons elicits food intake and operant water seeking (Zhu et al., 2017; Stephenson-Jones et al., 2020) and is reinforcing (Zhu et al., 2017; Faget et al., 2018), while optogenetic stimulation of VP glutamate neurons elicits aversive responses and promotes operant avoidance (Faget et al., 2018; Tooley et al., 2018; Levi et al., 2019; Stephenson-Jones et al., 2020)—though a recent report suggests that VP glutamate neurons may mediate salience irrespective of valence (Wang et al., 2020). None of these prior mouse studies indicated a role for VP^GABA^ neurons in aversive motivation, but they did not examine more complex types of aversive responding.

To address the function of VP^GABA^ neurons further, we examined the contribution of VP^GABA^ neurons to shock-induced affective responses, and to instrumental responding to avoid or escape shocks. Inhibiting VP^GABA^ neurons failed to impact shock-induced motor reactions or ultrasonic vocalizations, suggesting that these cells do not mediate aversion *per se*. However, when rats were trained to press a lever either to avoid an impending shock or to escape an ongoing one, DREADD inhibition revealed a hidden role for VP^GABA^ neurons in aversive motivation. Specifically, the latency to press a lever in order to cancel an impending shock was increased by VP^GABA^ inhibition, while latency to press to escape an ongoing shock was unaffected. Together, these data show that VP^GABA^ inhibition affected neither affective reactions to shock itself, nor the ability of an ongoing shock to induce escape responses. Instead, VP^GABA^-inhibited rats simply appeared to less urgently avoid impending punishment (though the proportion of trials escaped versus avoided was not altered). This increase in avoidance latency represents a departure from the common notion that VP^GABA^ neurons are solely implicated in appetitive behavior. Rather, these neurons seem instead to facilitate high-stakes instrumental behavior of many types. This said, we note that when rats pressed to avoid footshock, they also received a 20 sec signal indicating freedom from impending threat. It is therefore possible that DREADD inhibition did not impact aversive motivation itself, but instead reduced the conditioned reinforcing properties of this safety signal (Fernando et al., 2014). Dissociating avoidance of harm from pursuit of safety is famously difficult (LeDoux et al., 2017; Sangha et al., 2020), so further work is needed to disambiguate this newly-discovered role for VP^GABA^ neurons in aversive motivation.

### Specificity of Effects

We found very little evidence of non-selective effects of CNO in WT rats without DREADDs. In the absence of DREADDs, CNO can have off-target behavioral effects in some experiments (MacLaren et al., 2016; Gomez et al., 2017; Manvich et al., 2018). Yet across the numerous behaviors tested here, we identified only a trend towards an increase in escape latency in WT rats—indicating predominantly DREADD-specific effects of CNO (Mahler and Aston-Jones, 2018). In addition, although VP is sometimes considered a motor structure (Mogenson et al., 1980; Heimer et al., 1982), it is unlikely that VP^GABA^ DREADD effects were due to nonspecific motoric inhibition. Neither horizontal locomotion nor rearing behavior were affected by engaging VP^GABA^ DREADDs, and behavioral effects were specific to highly-motivated instrumental contexts—other behaviors like spontaneous chocolate intake, and pressing for chow in a sated state were unaffected.

VP DREADD expression was mostly localized within strictly-defined VP borders here, though in most rats at least some expression encroached upon nearby subcortical structures containing GABAergic neurons with important behavioral roles (Koob, 2004; Silberman et al., 2009; Jennings et al., 2013; Barker et al., 2017; Saga et al., 2017; Gordon-Fennell et al., 2020), so we cannot definitively exclude overlapping roles for these neurons in behavioral effects.

Similar to our prior findings (Farrell et al., 2019), we saw no evidence of sex-dependent behavioral effects of chemogenetic VP manipulation, though we also cannot exclude this possibility since studies were not powered to fully explore this variable.

We note that a portion of VP^GABA^ neurons also express other peptides such as parvalbumin and enkephalin, and such co-expression may have functional implications. For example, VP parvalbumin neurons, which consist of both GABAergic and glutamatergic neurons, are necessary for both alcohol seeking and depression-like behavior (Knowland et al., 2017; Prasad et al., 2020). Moreover, VP^GABA^ neurons that co-express enkephalin drive cue-induced cocaine seeking (Heinsbroek et al., 2019). A significant portion of these VP subpopulations also express GABA markers, though others are glutamatergic. Since VP glutamate and GABA neurons have dissociable roles in behavior (Faget et al., 2018; Tooley et al., 2018; Stephenson-Jones et al., 2020), further work is needed to parse the relative functional roles played by VP cells expressing GABA, glutamate, various co-expressed proteins, and combinations thereof.

## Conclusion

These results demonstrate an essential role for VP^GABA^ neurons in high-stakes motivated behavior—be it to pursue valued rewards, to avoid impending harm, or to make important decisions when motivations are mixed. We show for the first time that VP^GABA^ neurons’ role in motivation impacts decision making, since inhibiting these cells yields a conservative, risk-averse decision-making strategy rather than a simple decrease in all reward seeking. If successfully harnessed therapeutically, we speculate that suppressing VP^GABA^ neuron activity might be useful for treating addiction, or other disorders of maladaptive, risky decision making.

## Acknowledgements

We thank Erik Castillo and Christina Ruiz for technical assistance, Andrew M. Delamater for helpful comments on these data, and Ronald E. See for assistance with data collection.

## Notes

### Competing Interest Statement

The authors have declared no competing interest.

## References

Armbruster BN, Li X, Pausch MH, Herlitze S, Roth BL (2007) Evolving the lock to fit the key to create a family of G protein-coupled receptors potently activated by an inert ligand. Proceedings of the National Academy of Sciences 104:5163–5168.

Barker DJ, Miranda-Barrientos J, Zhang S, Root DH, Wang H-L, Liu B, Calipari ES, Morales M (2017) Lateral preoptic control of the lateral habenula through convergent glutamate and GABA transmission. Cell reports 21:1757–1769.

Berridge KC, Robinson TE (2003) Parsing reward. Trends in neurosciences 26:507–513.

Berridge KC, Kringelbach ML (2015) Pleasure systems in the brain. Neuron 86:646–664.

Bonnet KA, Peterson KE (1975) A modification of the jump-flinch technique for measuring pain sensitivity in rats. Pharmacology Biochemistry and Behavior 3:47–55.

Brudzynski SM (2013) Ethotransmission: communication of emotional states through ultrasonic vocalization in rats. Current opinion in neurobiology 23:310–317.

Castro DC, Cole SL, Berridge KC (2015) Lateral hypothalamus, nucleus accumbens, and ventral pallidum roles in eating and hunger: interactions between homeostatic and reward circuitry. Frontiers in systems neuroscience 9:90.

Chang SE, Todd TP, Bucci DJ, Smith KS (2015) Chemogenetic manipulation of ventral pallidal neurons impairs acquisition of sign-tracking in rats. European Journal of Neuroscience 42:3105–3116.

Chang SE, Smedley EB, Stansfield KJ, Stott JJ, Smith KS (2017) Optogenetic inhibition of ventral pallidum neurons impairs context-driven salt seeking. Journal of Neuroscience 37:5670–5680.

Creed M, Ntamati NR, Chandra R, Lobo MK, Lüscher C (2016) Convergence of reinforcing and anhedonic cocaine effects in the ventral pallidum. Neuron 92:214–226.

Cromwell HC, Berridge KC (1993) Where does damage lead to enhanced food aversion: the ventral pallidum/substantia innominata or lateral hypothalamus? Brain research 624:1–10.

Faget L, Zell V, Souter E, McPherson A, Ressler R, Gutierrez-Reed N, Yoo JH, Dulcis D, Hnasko TS (2018) Opponent control of behavioral reinforcement by inhibitory and excitatory projections from the ventral pallidum. Nature communications 9:849.

Farrar AM, Font L, Pereira M, Mingote S, Bunce JG, Chrobak JJ, Salamone JD (2008) Forebrain circuitry involved in effort-related choice: Injections of the GABAA agonist muscimol into ventral pallidum alter response allocation in food-seeking behavior. Neuroscience 152:321–330.

Farrell MR, Ruiz CM, Castillo E, Faget L, Khanbijian C, Liu S, Schoch H, Rojas G, Huerta MY, Hnasko TS (2019) Ventral pallidum is essential for cocaine relapse after voluntary abstinence in rats. Neuropsychopharmacology:1–13.

Fernando ABP, Urcelay GP, Mar AC, Dickinson A, Robbins TW (2014) Safety signals as instrumental reinforcers during free-operant avoidance. Learning & Memory 21:488–497.

Floresco SB, Braaksma DN, Phillips AG (1999) Involvement of the ventral pallidum in working memory tasks with or without a delay. Annals of the New York Academy of Sciences 877:711–716.

Floresco SB, Onge JRS, Ghods-Sharifi S, Winstanley CA (2008) Cortico-limbic-striatal circuits subserving different forms of cost-benefit decision making. Cognitive, Affective, & Behavioral Neuroscience 8:375–389.

Fujimoto A, Hori Y, Nagai Y, Kikuchi E, Oyama K, Suhara T, Minamimoto T (2019) Signaling incentive and drive in the primate ventral pallidum for motivational control of goal-directed action. Journal of Neuroscience 39:1793–1804.

Gallo EF, Meszaros J, Sherman JD, Chohan MO, Teboul E, Choi CS, Moore H, Javitch JA, Kellendonk C (2018) Accumbens dopamine D2 receptors increase motivation by decreasing inhibitory transmission to the ventral pallidum. Nature communications 9:1086.

Gibson GD, Prasad AA, Jean-Richard-dit-Bressel P, Yau JOY, Millan EZ, Liu Y, Campbell EJ, Lim J, Marchant NJ, Power JM (2018) Distinct accumbens shell output pathways promote versus prevent relapse to alcohol seeking. Neuron 98:512–520.

Gomez JL, Bonaventura J, Lesniak W, Mathews WB, Sysa-Shah P, Rodriguez LA, Ellis RJ, Richie CT, Harvey BK, Dannals RF (2017) Chemogenetics revealed: DREADD occupancy and activation via converted clozapine. Science 357:503–507.

Gordon-Fennell AG, Will RG, Ramachandra V, Gordon-Fennell L, Dominguez JM, Zahm DS, Marinelli M (2020) The lateral preoptic area: a novel regulator of reward seeking and neuronal activity in the ventral tegmental area. Frontiers in neuroscience 13:1433.

Heimer L, Switzer RD, Van Hoesen GW (1982) Ventral striatum and ventral pallidum: Components of the motor system? Trends in Neurosciences 5:83–87.

Heinsbroek J, Bobadilla A-C, Dereschewitz E, Assali A, Chalhoub RM, Cowan CW, Kalivas PW (2019) Opposing Regulation of Cocaine Seeking by Glutamate and Enkephalin Neurons in the Ventral Pallidum. CELL-REPORTS-D-19-03631.

Heinsbroek JA, Neuhofer DN, Griffin WC, Siegel GS, Bobadilla A-C, Kupchik YM, Kalivas PW (2017) Loss of plasticity in the D2-accumbens pallidal pathway promotes cocaine seeking. Journal of Neuroscience 37:757–767.

Jennings JH, Sparta DR, Stamatakis AM, Ung RL, Pleil KE, Kash TL, Stuber GD (2013) Distinct extended amygdala circuits for divergent motivational states. Nature 496:224–228.

Kelley AE, Baldo BA, Pratt WE, Will MJ (2005) Corticostriatal-hypothalamic circuitry and food motivation: integration of energy, action and reward. Physiology & behavior 86:773–795.

Knutson B, Burgdorf J, Panksepp J (2002) Ultrasonic vocalizations as indices of affective states in rats. Psychological bulletin 128:961.

Koob GF (2004) A role for GABA mechanisms in the motivational effects of alcohol. Biochemical pharmacology 68:1515–1525.

Kupchik YM, Kalivas PW (2013) The rostral subcommissural ventral pallidum is a mix of ventral pallidal neurons and neurons from adjacent areas: an electrophysiological study. Brain Structure and Function 218:1487–1500.

Kupchik YM, Brown RM, Heinsbroek JA, Lobo MK, Schwartz DJ, Kalivas PW (2015) Coding the direct/indirect pathways by D1 and D2 receptors is not valid for accumbens projections. Nature neuroscience 18:1230.

LeDoux JE, Moscarello J, Sears R, Campese V (2017) The birth, death and resurrection of avoidance: a reconceptualization of a troubled paradigm. Molecular psychiatry 22:24–36.

Levi LA, Inbar K, Nachshon N, Bernat N, Gatterer A, Inbar D, Kupchik YM (2019) Projection-specific potentiation of ventral pallidal glutamatergic outputs after abstinence from cocaine. Journal of Neuroscience.

MacLaren DAA, Browne RW, Shaw JK, Radhakrishnan SK, Khare P, España RA, Clark SD (2016) Clozapine N-oxide administration produces behavioral effects in Long–Evans rats: implications for designing DREADD experiments. eneuro 3.

Mahler SV, Aston-Jones G (2018) CNO Evil? Considerations for the use of DREADDs in behavioral neuroscience. Neuropsychopharmacology 43:934.

Mahler SV, Smith RJ, Aston-Jones G (2013a) Interactions between VTA orexin and glutamate in cue-induced reinstatement of cocaine seeking in rats. Psychopharmacology 226:687–698.

Mahler SV, Moorman DE, Feltenstein MW, Cox BM, Ogburn KB, Bachar M, McGonigal JT, Ghee SM, See RE (2013b) A rodent “self-report” measure of methamphetamine craving? Rat ultrasonic vocalizations during methamphetamine self-administration, extinction, and reinstatement. Behavioural brain research 236:78–89.

Mahler SV, Vazey EM, Beckley JT, Keistler CR, McGlinchey EM, Kaufling J, Wilson SP, Deisseroth K, Woodward JJ, Aston-Jones G (2014) Designer receptors show role for ventral pallidum input to ventral tegmental area in cocaine seeking. Nature neuroscience 17:577.

Mahler SV, Brodnik ZD, Cox BM, Buchta WC, Bentzley BS, Quintanilla J, Cope ZA, Lin EC, Riedy MD, Scofield MD (2019) Chemogenetic Manipulations of Ventral Tegmental Area Dopamine Neurons Reveal Multifaceted Roles in Cocaine Abuse. Journal of Neuroscience 39:503–518.

Manvich DF, Webster KA, Foster SL, Farrell MS, Ritchie JC, Porter JH, Weinshenker D (2018) The DREADD agonist clozapine N-oxide (CNO) is reverse-metabolized to clozapine and produces clozapine-like interoceptive stimulus effects in rats and mice. Scientific reports 8:3840.

McAlonan GM, Robbins TW, Everitt BJ (1993) Effects of medial dorsal thalamic and ventral pallidal lesions on the acquisition of a conditioned place preference: further evidence for the involvement of the ventral striatopallidal system in reward-related processes. Neuroscience 52:605–620.

Meye FJ, Soiza-Reilly M, Smit T, Diana MA, Schwarz MK, Mameli M (2016) Shifted pallidal co-release of GABA and glutamate in habenula drives cocaine withdrawal and relapse. Nature neuroscience 19:1019–1024.

Mitchell MR, Weiss VG, Beas BS, Morgan D, Bizon JL, Setlow B (2014) Adolescent risk taking, cocaine self-administration, and striatal dopamine signaling. Neuropsychopharmacology 39:955.

Mogenson GJ, Jones DL, Yim CY (1980) From motivation to action: functional interface between the limbic system and the motor system. Progress in neurobiology 14:69–97.

Morgane PJ (1961) Alterations in feeding and drinking behavior of rats with lesions in globi pallidi. American Journal of Physiology-Legacy Content 201:420–428.

Oleson EB, Gentry RN, Chioma VC, Cheer JF (2012) Subsecond dopamine release in the nucleus accumbens predicts conditioned punishment and its successful avoidance. Journal of Neuroscience 32:14804–14808.

Onge JRS, Abhari H, Floresco SB (2011) Dissociable contributions by prefrontal D1 and D2 receptors to risk-based decision making. Journal of Neuroscience 31:8625–8633.

Orsini CA, Trotta RT, Bizon JL, Setlow B (2015a) Dissociable roles for the basolateral amygdala and orbitofrontal cortex in decision-making under risk of punishment. Journal of Neuroscience 35:1368–1379.

Orsini CA, Moorman DE, Young JW, Setlow B, Floresco SB (2015b) Neural mechanisms regulating different forms of risk-related decision-making: Insights from animal models. Neuroscience & Biobehavioral Reviews 58:147–167.

Orsini CA, Willis ML, Gilbert RJ, Bizon JL, Setlow B (2016) Sex differences in a rat model of risky decision making. Behavioral neuroscience 130:50.

Orsini CA, Hernandez CM, Singhal S, Kelly KB, Frazier CJ, Bizon JL, Setlow B (2017) Optogenetic inhibition reveals distinct roles for basolateral amygdala activity at discrete time points during risky decision making. Journal of Neuroscience 37:11537–11548.

Ottenheimer D, Richard JM, Janak PH (2018) Ventral pallidum encodes relative reward value earlier and more robustly than nucleus accumbens. Nature communications 9:4350.

O’Neal TJ, Nooney MN, Thien K, Ferguson SM (2019) Chemogenetic modulation of accumbens direct or indirect pathways bidirectionally alters reinstatement of heroin-seeking in high-but not low-risk rats. Neuropsychopharmacology:1–12.

Paxinos G, Watson C (2006) The rat brain in stereotaxic coordinates: hard cover edition: Elsevier.

Pessiglione M, Schmidt L, Draganski B, Kalisch R, Lau H, Dolan RJ, Frith CD (2007) How the brain translates money into force: a neuroimaging study of subliminal motivation. Science 316:904–906.

Pleil KE, Rinker JA, Lowery-Gionta EG, Mazzone CM, McCall NM, Kendra AM, Olson DP, Lowell BB, Grant KA, Thiele TE (2015) NPY signaling inhibits extended amygdala CRF neurons to suppress binge alcohol drinking. Nature neuroscience 18:545–552.

Portfors CV (2007) Types and functions of ultrasonic vocalizations in laboratory rats and mice. Journal of the American Association for Laboratory Animal Science 46:28–34.

Prasad AA, McNally GP (2020) The ventral pallidum and relapse to alcohol seeking. British Journal of Pharmacology.

Richard JM, Ambroggi F, Janak PH, Fields HL (2016) Ventral pallidum neurons encode incentive value and promote cue-elicited instrumental actions. Neuron 90:1165–1173.

Rogers JL, Ghee S, See RE (2008) The neural circuitry underlying reinstatement of heroin-seeking behavior in an animal model of relapse. Neuroscience 151:579–588.

Root DH, Melendez RI, Zaborszky L, Napier TC (2015) The ventral pallidum: Subregion-specific functional anatomy and roles in motivated behaviors. Progress in neurobiology 130:29–70.

Roth BL (2016) DREADDs for neuroscientists. Neuron 89:683–694.

Saga Y, Hoshi E, Tremblay L (2017) Roles of multiple globus pallidus territories of monkeys and humans in motivation, cognition and action: an anatomical, physiological and pathophysiological review. Frontiers in neuroanatomy 11:30.

Saga Y, Richard A, Sgambato-Faure V, Hoshi E, Tobler PN, Tremblay L (2016) Ventral pallidum encodes contextual information and controls aversive behaviors. Cerebral Cortex 27:2528–2543.

Salamone JD, Correa M (2012) The mysterious motivational functions of mesolimbic dopamine. Neuron 76:470–485.

Sangha S, Diehl MM, Bergstrom HC, Drew MR (2020) Know safety, no fear. Neuroscience & Biobehavioral Reviews 108:218–230.

Sharpe MJ, Marchant NJ, Whitaker LR, Richie CT, Zhang YJ, Campbell EJ, Koivula PP, Necarsulmer JC, Mejias-Aponte C, Morales M (2017) Lateral hypothalamic GABAergic neurons encode reward predictions that are relayed to the ventral tegmental area to regulate learning. Current Biology 27:2089–2100.

Silberman Y, Bajo M, Chappell AM, Christian DT, Cruz M, Diaz MR, Kash T, Lack AK, Messing RO, Siggins GR (2009) Neurobiological mechanisms contributing to alcohol–stress–anxiety interactions. Alcohol 43:509–519.

Simmons WK, Rapuano KM, Ingeholm JE, Avery J, Kallman S, Hall KD, Martin A (2014) The ventral pallidum and orbitofrontal cortex support food pleasantness inferences. Brain Structure and Function 219:473–483.

Simon NW, Setlow B (2012) Modeling risky decision making in rodents. In: Psychiatric Disorders, pp 165–175: Springer.

Simon NW, Gilbert RJ, Mayse JD, Bizon JL, Setlow B (2009) Balancing risk and reward: a rat model of risky decision making. Neuropsychopharmacology 34:2208.

Smith KS, Berridge KC (2005) The ventral pallidum and hedonic reward: neurochemical maps of sucrose “liking” and food intake. Journal of neuroscience 25:8637–8649.

Smith KS, Berridge KC (2007) Opioid limbic circuit for reward: interaction between hedonic hotspots of nucleus accumbens and ventral pallidum. Journal of neuroscience 27:1594–1605.

Smith KS, Tindell AJ, Aldridge JW, Berridge KC (2009) Ventral pallidum roles in reward and motivation. Behavioural brain research 196:155–167.

Smith RJ, Aston-Jones G (2012) Orexin/hypocretin 1 receptor antagonist reduces heroin self-administration and cue-induced heroin seeking. European Journal of Neuroscience 35:798–804.

Spear LP (2000) The adolescent brain and age-related behavioral manifestations. Neuroscience & biobehavioral reviews 24:417–463.

Stephenson-Jones M, Bravo-Rivera C, Ahrens S, Furlan A, Xiao X, Fernandes-Henriques C, Li B (2020) Opposing contributions of GABAergic and glutamatergic ventral pallidal neurons to motivational behaviors. Neuron 105:921–933.

Stratford TR, Kelley AE, Simansky KJ (1999) Blockade of GABAA receptors in the medial ventral pallidum elicits feeding in satiated rats. Brain research 825:199–203.

Tachibana Y, Hikosaka O (2012) The primate ventral pallidum encodes expected reward value and regulates motor action. Neuron 76:826–837.

Tindell AJ, Smith KS, Berridge KC, Aldridge JW (2009) Dynamic computation of incentive salience:”wanting” what was never “liked”. Journal of Neuroscience 29:12220–12228.

Tooley J, Marconi L, Alipio JB, Matikainen-Ankney B, Georgiou P, Kravitz AV, Creed MC (2018) Glutamatergic ventral pallidal neurons modulate activity of the habenula–tegmental circuitry and constrain reward seeking. Biological psychiatry 83:1012–1023.

Turner MS, Gray TS, Mickiewicz AL, Napier TC (2008) Fos expression following activation of the ventral pallidum in normal rats and in a model of Parkinson’s Disease: implications for limbic system and basal ganglia interactions. Brain Structure and Function 213:197–213.

Wakabayashi KT, Feja M, Baindur AN, Bruno MJ, Bhimani RV, Park J, Hausknecht K, Shen R-Y, Haj-Dahmane S, Bass CE (2019) Chemogenetic activation of ventral tegmental area GABA neurons, but not mesoaccumbal GABA terminals, disrupts responding to reward-predictive cues. Neuropsychopharmacology 44:372.

Wang F, Zhang J, Yuan Y, Chen M, Gao Z, Zhan S, Fan C, Sun W, Hu J (2020) Salience processing by glutamatergic neurons in the ventral pallidum. Science Bulletin 65:389–401.

Zhu C, Yao Y, Xiong Y, Cheng M, Chen J, Zhao R, Liao F, Shi R, Song S (2017) Somatostatin neurons in the basal forebrain promote high-calorie food intake. Cell reports 20:112–123.

